# Yield maintenance under drought is orchestrated by the *qDTY12.1*-encoded *DECUSSATE* gene of rice through a network with other flowering-associated genes across the genetic background

**DOI:** 10.1101/2021.02.09.430414

**Authors:** Jacobo Sanchez, Pushpinder Pal Kaur, Isaiah C.M. Pabuayon, Naga Bhushana Rao Karampudi, Ai Kitazumi, Nitika Sandhu, Margaret Catolos, Arvind Kumar, Benildo G. de los Reyes

**Affiliations:** Department of Plant and Soil Science, Texas Tech University, Lubbock, USA; International Rice Research Institute, Los Banos, Philippines

## Abstract

Introgression of major-effect QTLs is an important component of rice breeding for yield-retention under drought. While largely effective, the maximum potentials of such QTLs have not been consistent across genetic backgrounds. We hypothesized that synergism or antagonism with additive-effect peripheral genes across the background could either enhance or undermine the QTL effects. To elucidate the molecular underpinnings of such interaction, we dissected *qDTY12.1* synergy with numerous peripheral genes in context of network rewiring effects. By integrative transcriptome profiling and network modeling, we identified the *DECUSSATE* (*OsDEC*) within *qDTY12.1* as the core of the synergy and shared by two sibling introgression lines in IR64 genetic background, *i.e.,* LPB (low-yield penalty) and HPB (high-yield penalty). *OsDEC* is expressed in flag leaves and induced by progressive drought at booting stage in LPB but not in HPB. The unique *OsDEC* signature in LPB is coordinated with 35 upstream and downstream peripheral genes involved in floral development through the cytokinin signaling pathway, which are lacking in HPB. Results further support the differential network rewiring effects through genetic coupling-uncoupling between *qDTY12.1* and other upstream and downstream peripheral genes across the distinct genetic backgrounds of LPB and HPB. We propose that the functional *DEC*-network in LPB defines a mechanism for early flowering as a means for avoiding the depletion of photosyntate needed for reproductive growth due to drought. Its impact on yield-retention is likely through the timely establishment of stronger source-sink dynamics that sustains a robust reproductive transition under drought.

**Author summary:** While the Green Revolution of the 1960’s significantly increased rice grain yields through the creation of high-yielding varieties for high input systems, current marginal climates pose a significant challenge for providing consistent yield. In rice growing regions of the world, drought affects the livelihood of small-scale and subsistence farmers by inflicting significant yield penalties to their production systems. Breeding of next-generation rice varieties with optimal balance of survivability and productivity traits will be key to providing consistent yields year to year. Within this paradigm, the use of large effect QTLs such as *qDTY12.1* to improve yield retention under drought have been largely successful. By integrating the use of high resolution transcriptome datasets with a focused biological interrogation of agronomic results from this and previous studies, we uncovered a putative functional genetic network, anchored by the *DECUSSATE* gene (*OsDEC*) within *qDTY12.1*, that effectively minimizes drought penalties to yield by driving cellular processes that culminate in timely flowering that maximizes the use of photosynthetic sources for efficient reproduductive transition and ultimately seed development. Our study further illuminates the *qDTY12.1* function and speaks to the misconception that *qDTY* introgression alone is sufficient for providing consistently large positive effects to yield retention under reproductive stage drought.

## Introduction

Through ideotype breeding, the Green Revolution created the modern high-yielding varieties of rice with morphological and physiological attributes optimal for environments with ample water and nutrients **[1–6]**. However, the increased incidence of erratic rainfall patterns, diminishing water resources, and depletion of arable lands paints the new reality at which crop production must be undertaken to ensure yield stability under increasing global food demands and steadily rising population. With this reality in mind, innovative and holistic paradigms in plant breeding will be critical to the development of the next generation of crop cultivars that can absorb this conglomeration of ecological factors while minimizing penalties to yield. The creation of new ideotypes with novel mechanisms that confer resilience to drought-prone environments for example, holds great promise for establishing an effective means for maximizing yield under sub-optimal conditions **[7,8]**.

With the reported drought-related yield losses in rice ranging from 18% to 97% **[9]**, robust approaches in breeding, like QTL introgression and pyramiding for example, become the most vital components of a holistic strategy for addressing the needs of subsistence rice farmers in regions that are highly prone to either periodic episodes of drought or persistent drought **[10–13]**. The discovery and subsequent pyramiding of large-effect QTLs that function at the reproductive stage, *i.e., qDTYs* for yield maintenance under drought **(S1 Table)**, have led to incremental but major improvements in the yield potential of many of the widely grown rice cultivars that regularly incur significant penalties due to reproductive-stage drought **[14–16]**. Among the most well-characterized and considered of very high importance to rice breeding is the *qDTY12.1*, because of its more consistent effects in reducing the penalty to yield across growing environments **[17–19]**. Fine-mapping of *qDTY12.1* in the Way Rarem x Vandana derived population defined its boundaries within 3.1cM on the long-arm of chromosome-12, which is estimated to be about 1.554 Mbp with physical coordinates in the Nipponbare *RefSeq* between 15,848,736 bp to 17,401,530 bp **[20]**.

Initial attempts to understand the mechanisms by which *qDTY12.1* is able to impart such large positive effects as a ‘*yield QTL’* have pointed to a number of candidate genes **[21–23]**. However, while the characterization of these genes provided important advances, much of the mechanisms that have been uncovered so far appeared to be involved in stress avoidance and physiological adjustments during vegetative growth, and not really in the cellular processes with direct significance to reproductive growth, source-sink partitioning, and/or grain development, which are more meaningful to yield maintenance **[24–27]**. For instance, a recent study showed the importance of the *qDTY12.1*-encoded *OsNAM_12.1_* transcription factor *(Os12g0477400*) in the regulation of root development and architecture as a mechanism of drought avoidance during vegetative growth **[23]**. Additionally, a meta-analysis of 53 grain yield-related QTLs identified six (6) loci within the meta-QTL (MQTL) on *qDTY12.1* that are not directly associated with yield processes **[21]**.

Perhaps the most interesting aspect of *qDTY12.1* was the fact that this locus did not exhibit a positive effect on yield maintenance in its native genetic background (*i.e.,* original donor), which is the Indonesian upland cultivar Way Rarem (WR) **[17]**. However, significant positive effects of the *qDTY12.1* in minimizing yield penalty under drought were observed in recombinants with the Indian cultivar Vandana, which has drought tolerance at the vegetative stage but with high drought penalty to yield **[17,28]**. These seminal observations inspired the initial hypothesis that the full effects of *qDTY12.1* require some kind of synergy and complementation with other minor peripheral genes in the genetic background that cannot be identified at high statistical confidence by the resolution of QTL mapping **[22]**. Researchers have been trying to identify such network of genes either among the *qDTY12.1* genes themselves or across genetic backgrounds, but so far no truly significant leads apart from vegetative stage drought avoidance have been uncovered **[24–27]**.

With the observations that some introgression derivatives of *qDTY12.1* exhibited consistent yield retention under drought, while others had not, the question was raised as to why the presence of the *qDTY12.1* allele of WR alone as facilitated by marker-assisted selection, would not be sufficient in providing the expected positive effects across different genetic backgrounds or even within similar genetic backgrounds **[22]**. We hypothesized that in specific derivatives carrying the same *qDTY12.1* allele from WR where the expected positive effects were not manifested, genetic recombination may have created some kind of coupling-uncoupling effects involving many other alleles in the genetic background that are peripheral but synergistic to *qDTY12.1* functions, with the *qDTY12.1* genes themselves acting as the core of the mechanism (*i.e.,* epistatic effects or network rewiring effects) **[29,30]**. This hypothesis built its strength from the recently proposed *Omnigenic Theory,* which postulated that complex traits are controlled by not only a core set of loci with quantifiable effects, but also by a genome-wide cohort of other peripheral loci whose individual effects are minute but their additive effects could either positively or negatively complement the core effects to account for a larger proportion of the total phenotypic variance **[31]**.

To dig deeper into the yield-related function of *qDTY12.1* while also addressing the coupling-uncoupling and network rewiring hypotheses, we investigated a minimal comparative panel established at the International Rice Research Institute (IRRI) that models the contrast between positive net gain and negative net gain from *qDTY12.1* effects across potentially contrasting combinations of peripheral alleles in similar genetic backgrounds **[32]**. This comparative panel is comprised of the cultivar Way Rarem (WR), the original donor of *qDTY12.1*, the drought-sensitive mega-variety IR64 as the recipient of *qDTY12.1* from WR, and two IR64 sibling backcross derivatives with the *qDTY12.1* of WR introgressed through a bridge donor recombinant with Vandana, hence *Low Yield Penalty* (LPB) and *High Yield Penalty* (HPB) introgression lines **(S1 Fig) [9]**.

A cautionary thinking is that LPB and HPB are considered to have uniform genetic backgrounds but only at the extent and resolution afforded by marker-based genotyping, and not based on whole-genome sequence assembly. That being said, the potential contributions of other hidden introgressions that could possibly be traced from other donors in their pedigrees (*i.e.,* either WR or Vandana), beyond what can be ascertained by the resolution of marker-assisted selection of the foreground and background, must not be excluded as potential sources of cryptic variations between LPB and HPB. By in-depth analysis of the drought-response transcriptomes at vegetative, reproductive (booting), and grain filling stages under field drought conditions, along with the modeling of co-expression networks, we identified the first candidate gene of *qDTY12.1* with a convincing direct link to processes that may modulate the timing of reproductive transition under the limiting source-sink status during drought. We report here the identification of *DECUSSATE* gene *(OsDEC)*, a single copy locus in the rice genome (Os12g0465700) and first identified as a regulator of leaf phyllotaxy **[33]**, as a crucial gene of *qDTY12.1* that facilitates efficient panicle development under drought, mediated by cytokinin.

## Results

### Agronomic performances under drought across the comparative panel

The comparative panel was subjected to slow but progressive drought in the rain-sheltered drought facility at IRRI from the mid-vegetative stage through the grain-filling stage **(S2 Fig) [34]**. Integrative analysis of grain yield data extracted from previously published studies under identical drought experimental conditions at IRRI **[22]**, revealed that LPB suffered a 74% yield penalty from drought, while HPB, WR, and IR64 suffered higher yield penalties of 97.5%, 94.6%, and 89.1%, respectively (Fig. 1A). Analysis of yield data from identical field drought experiments performed at IRRI for this transcriptome study recapitulated the same trends, with 66.3% penalty for LPB, 87.1% for HPB, and 77.3% for IR64 **[22]**. However, WR showed a lower penalty (58.3%) than previously observed (Fig 1B). While there were year-to-year variations, it was evident that LPB consistently outperformed the other genotypes.

**Fig 1.**
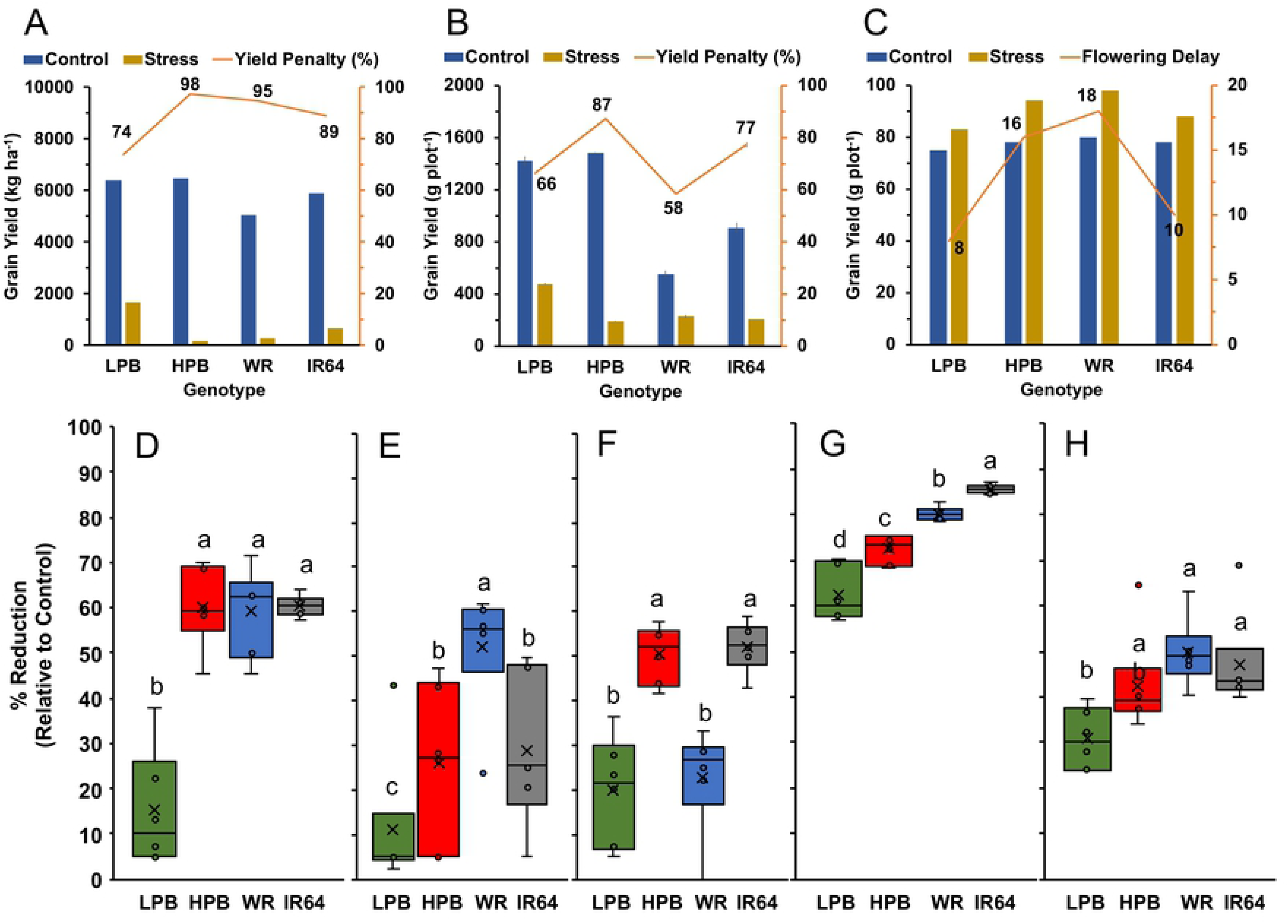
Synthesis and integration of all available data on relative agronomic and yield performances across the minimal comparative panel during progressive drought. Data from previous years of agronomic trials were integrated with the data collected from the 2017 wet season experiment performed for transcriptome studies. (**A**) Published grain yield results (GY; kg ha^-1^) **[22]** had significantly lower drought-mediated yield penalty (orange line) in LPB compared to the other genotypes. (**B**) Grain yield (g plot^-1^, n = 3, means ± s.e.) from the wet season-2017 experiment recapitulated the trends in previous years (orange line). (**C**) Drought-induced flowering delay (orange line) from published results **[22]** was also much less severe in LPB compared to the other genotypes. Trends in the (**D**) number of reproductive tillers per plant, (**E**) panicle length, (**F**) number of tillers per plant, (**G**) biomass per plant, and (**H**) plant height reiterated the superiority of LPB. Significant differences in flowering delay coupled with yield component reduction implied the earlier formation of reproductive sinks under drought in LPB, thus reducing grain yield penalty. LPB – Low Penalty BIL; HPB – High Penalty BIL; WR – Way Rarem (*qDTY12.1* donor); IR64 – rice mega variety (recurrent parent). Box plots with similar letters are not statistically significant at p < 0.05 using Tukey HSD (n=6).

Days to flowering also varied significantly across the comparative panel **[22]**. LPB showed a drought-induced delay in flowering of only 8 days compared to HPB, WR, and IR64, with delays of 16, 18, and 10 days, respectively (Fig 1C). These differences suggested that LPB may have established a stronger reproductive sink much earlier than the inferior genotypes, and this may have allowed an escape from the negative impacts of drought to resource allocation during the critical stages of floral organ development. Indeed, trends in five other growth components with direct significance to yield potential showed that LPB was superior with regard to the magnitude of drought-induced reductions in the number of reproductive tillers, panicle length, total number of tillers, dry biomass per plant, and plant height (Fig 1D **to 1H)**. Taken together, significant differences in grain yield and other agronomic attributes relevant to yield between LPB and HPB, suggest that while the sibling *qDTY12.1* introgression lines may be sharing largely similar genetic backgrounds, their yield potentials under drought were significantly different from each other.

### Transcriptome fluxes across genotypes revealed by Propensity normalization

Based on contrasting drought phenotypes, we hypothesized that fine-scale differences at the transcriptome level could be detected between LPB and HPB. Temporal fluxes in the transcriptome are windows to both subtle and large-scale differences between the sibling introgression lines that could illuminate potential differences in global regulatory mechanisms. Using the Propensity-normalized FPKM values, we performed two comparisons to capture the profiles of transcriptome fluxes in the flag leaves across genotypes and developmental stages. The first comparison utilized unfiltered Propensity-scores, *i.e.,* total distribution (*-n < Propensity > +n*) within three windows of the flag leaf transcriptomes, namely, global or total gene set (n = 25,786), and transcription factor (n = 1,340) and stress-related (n = 2,589) gene subsets **(S3 Fig; S2 Table)**. Hierarchical clustering indicated that in all three windows, the booting stage profiles exhibited significant dissimilarities between the four genotypes irrespective of growth condition. In contrast, LPB and HPB had very similar profiles at the vegetative stage under both irrigated and drought conditions with surprising similarity to WR, and dissimilarity to IR64 under irrigated condition. Fluxes during grain-filling showed significant overlaps across genotypes, but with LPB showing higher intensity in the positive propensity bands (Fig 2A). The similarities in vegetative profiles across LPB, HPB, and WR coupled with dissimilarity to IR64 was unexpected as the genomic contribution from WR was supposed to have been significantly diluted during recombination with Vandana and during the subsequent introgression to IR64 **[22]**.

**Fig 2.**
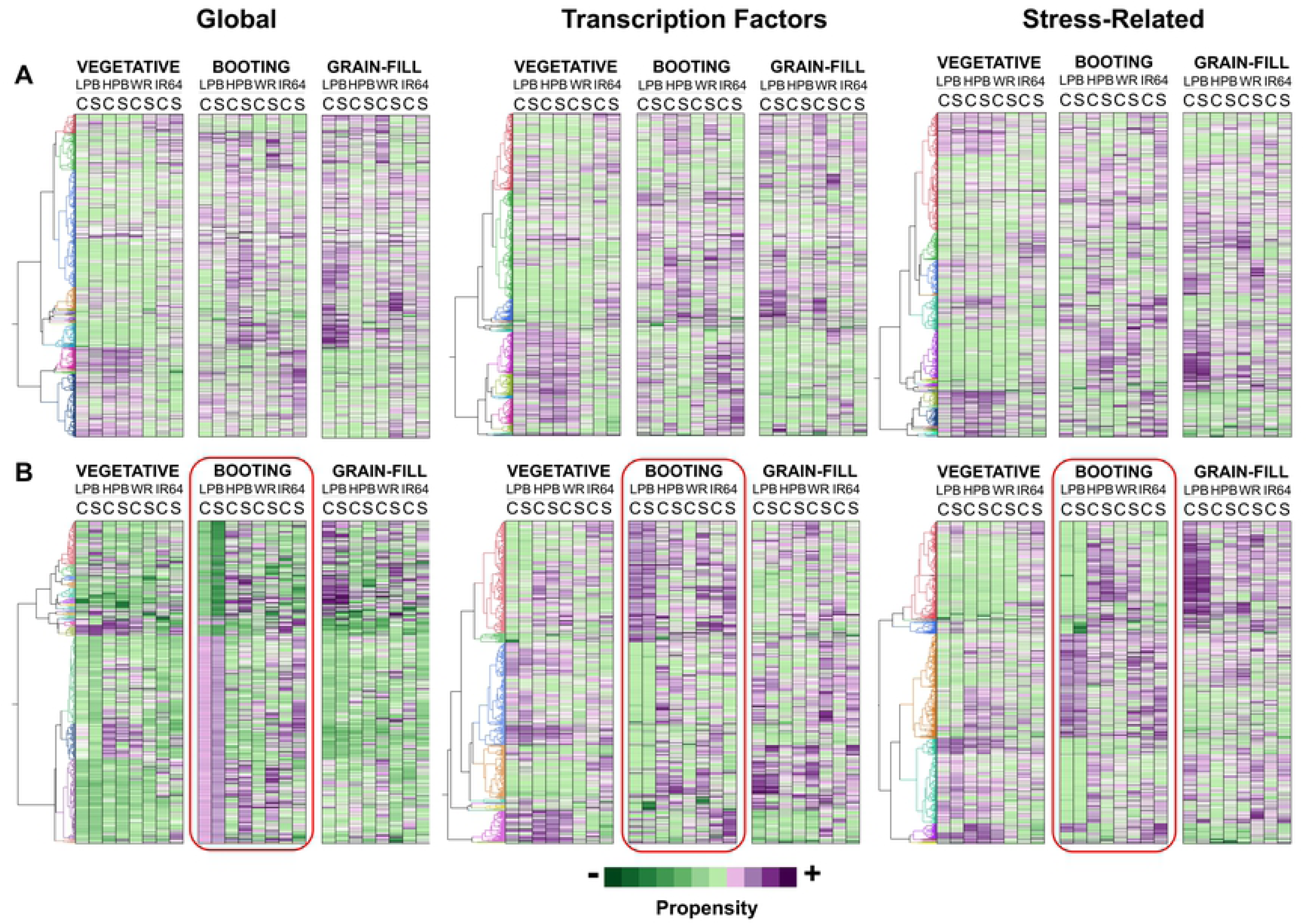
General trends in the transcriptomic fluxes across genotypes revealed by the filtered and un-filtered Propensity values of the global, transcription factor, and stress-related windows of the transcriptomes. (**A**) Hierarchical clustering of unfiltered global (27,786 loci), transcription factor (1,340 loci) and stress-related (2,589 loci) datasets. Expression fluxes at the vegetative stage under irrigated conditions highlight similarities between siblings (LPB, HPB) and WR but not IR64. Expression fluxes at the booting stage revealed the uniqueness of LPB, while grain-filling stage fluxes revealed high similarities across all genotypes. (**B**) Filtered (*-0.3 ≤ Propensity ≥+0.3*) transcriptome datasets included 384 (global), 410 (transcription factor), and 833 (stress-related) gene loci. This comparison recapitulated the general trends in the unfiltered datasets and further underscored the uniqueness of LPB, particularly during booting (red boxes). On a locus-by-locus comparison, during booting stage in LPB appeared to be well conserved between irrigated and drought. Fluxes in HPB, WR, and IR64 reflected a state of perturbation.

The second comparison was based on filtered Propensity scores of the global, transcription factor, and stress-related windows of the flag leaf transcriptomes. Genes included in this comparison had Propensity scores within the defined ranges of *-0.3 ≤ Propensity ≥ +0.3,* where the highest probabilities for significant differences in both positive and negative directions would be expected **(S3 Fig)**. Changes in expression among these genes were not due to spurious fluctuations and included 8,215 genes, 410 genes, and 833 genes in the global, transcription factor, and stress-related windows, respectively. Hierarchical clustering more vividly demonstrated the uniqueness of the fluxes of LPB at booting stage across the three windows (red boxes), and recapitulated the similarities between LPB, HPB, and WR at the vegetative stage under irrigated conditions, the dissimilarity with IR64 at vegetative stage, and the overlaps between the four genotypes at grain-filling stage (Fig 2B).

Conclusively, the LPB fluxes at booting stage showed a well-modulated character under both irrigated and drought conditions for the vast majority of genes across all three windows. In stark contrast, HPB, WR, and IR64 showed significant fragmentation of fluxes, giving evidence of a disjointed expression character. Altogether, the patterns revealed by both the filtered and unfiltered comparisons of Propensity-normalized expression established the uniqueness of LPB at booting stage, especially relative to its sibling HPB.

### Directionality of transcriptome fluxes suggests a robust mechanism in LPB

Integral to adaptive responses at the cellular level, the directional character (*i.e.,* upward skew, downward skew) of transcriptomic fluxes would be indicative of how well the complex waves of signals and gene activation and repression are organized in accordance with the underlying genetic circuitry towards cellular efficiency **[35]**. In conjunction with the flux analysis, we also determined the fraction of genes in the three flag leaf transcriptome windows with positive (positive propensity fraction; PPF) and negative (negative propensity fraction; NPF) propensity scores, respectively. These genes were correlated with the magnitude of skewing across the Propensity distributions across developmental stages in all three transcriptome windows (Fig 3**; S3 Fig**).

**Fig 3.**
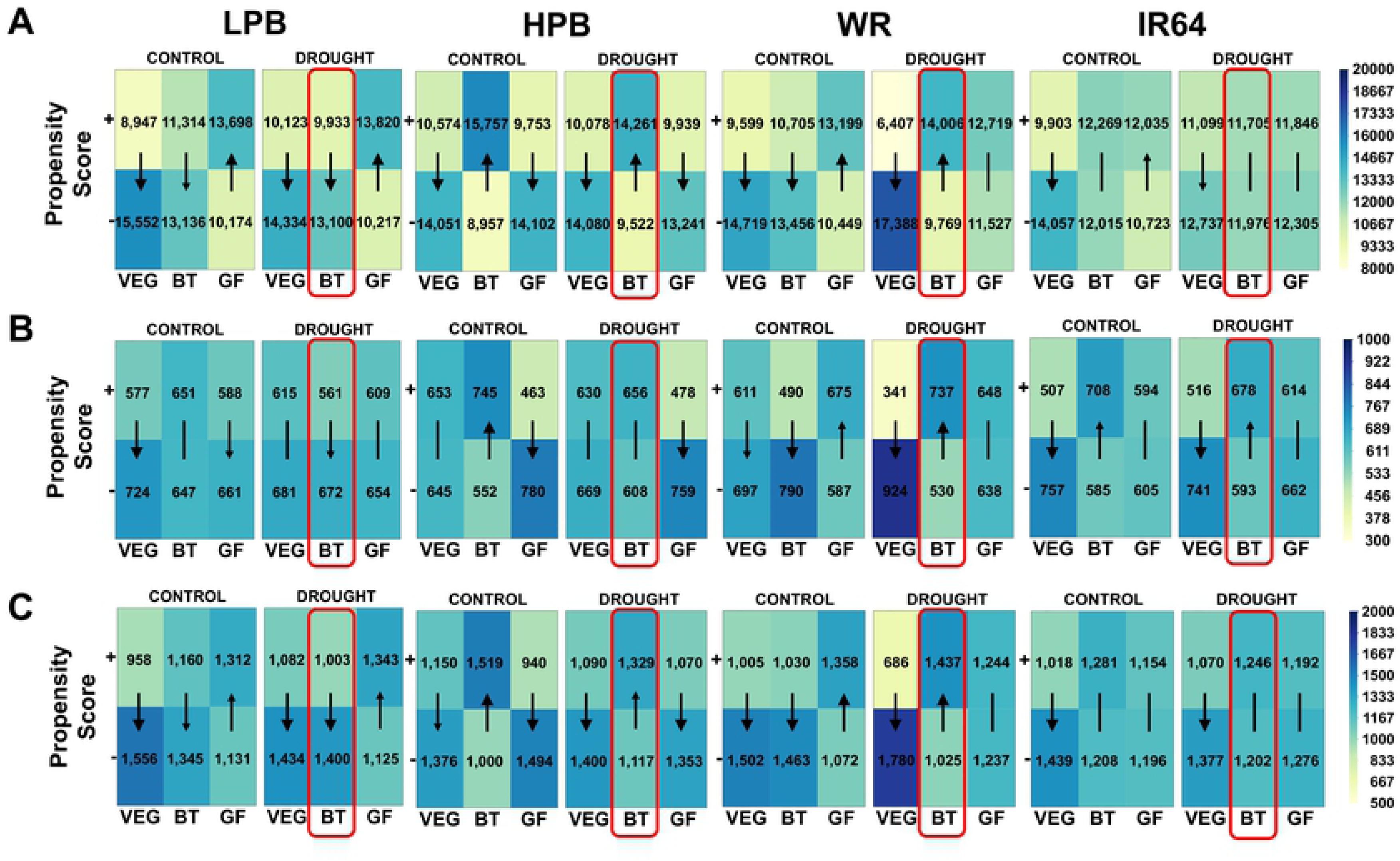
Differences in the directional character of transcriptomic fluxes across genotypes. Propensity scores of (**A**) global (25,786 loci), (**B**) transcription factor (1,340 loci), and (**C**) stress-related (2,589 loci) windows were divided into positive propensity (PPF) and negative propensity (NPF) fractions, excluding Propensity = 0. Directional characters were positive skew (upward arrow; PPF>NPF), negative skew (downward arrow; NPF>PPF) or neutral (line without arrow). Global, transcription factor, and stress-related windows showed negative skew in LPB, positive skew in HPB and WR, and neutral in IR64 at booting stage under drought (red boxes). The downward directional character of the LPB transcriptome at booting under drought illustrated a ‘*tamed*’ transcriptional landscape. Upward directional character in HPB and WR alluded to an ‘*untamed*’ or noisy transcriptional landscape.

Consistent with the unique fluxes observed in LPB at booting stage, the directional character of the three transcriptome windows was also unique in LPB at booting stage under drought (**red boxes in** Fig 3A**-3C)**, with LPB exhibiting a downward skew (NPF>PPF) whereas HPB and WR had upward skews (PPF>NPF), and IR64 being neutral. With very few exceptions, the directional character of each gene set was highly conserved between the irrigated and drought conditions within a genotype, irrespective of developmental stage. This potentially ‘*hard wired*’ nature of the directional character signified that expression fluxes that correlate with either positive or negative phenotype may have resulted from fine-scale dynamics of transcriptional modulation within specific subsets of genes (*i.e.,* networks). The downward directional character of LPB at booting stage during drought is an evidence of a ‘*tamed*’ transcriptome, where fluxes are highly organized and targeted for effective use of the transcriptional machinery without much trade-offs. In contrast, HPB and WR exhibited an ‘*untamed*’ hence highly active transcriptomes, betraying a disordered response with potentially detrimental consequences stemming from inefficient use of cellular resources **[36,37]**.

Booting stage represents a critical shift in resource allocation from vegetative sources to reproductive sinks that could be impaired drastically by drought **[38,39]**. Conservation and efficient use of cellular resources as mediated by the downward transcriptomic fluxes in LPB would prove beneficial for successful reproductive development. The unique signature towards more modulated fluxes in LPB implies an efficient resource allocation that may be impacting source-sink strength towards reproductive transition. This finding appears to be consistent with the function of *qDTY12.1* as a yield QTL **(S2 Fig)**.

Coupled with the downward, conservative fluxes at booting stage, LPB exhibited a positive skew across all three windows at the grain-filling stage during drought (Fig 3A**-3C)**. This was in contrast to HPB which showed a negative skew in the same three windows. Both WR and IR64 showed mostly non-skewed fluxes for all three windows under drought. The grain-filling stage represents the temporal continuum when the grain biomass is largely dependent on how well resources are channeled to reproductive sinks during development. Thus, the upward transcriptomic fluxes in LPB compared to the downward fluxes in HPB during the grain-filling stage may have contributed to their differences in yield retention. The directional character of the transcriptomic fluxes did not differ between the genotypes at the vegetative stage. However, there were differences in the magnitude of the skew with WR exhibiting the most drastic downward flux (Fig 3A**-3C)**.

Evidence for the directionality trends was also apparent in the Propensity distribution plots of the global transcriptomes across genotypes under both irrigated and drought conditions **(S3 Fig)**. The distribution plots at the vegetative stage under irrigated condition had almost direct overlap across all genotypes, signifying that the transcriptomes were all in homeostatic, low-level conditions (*i.e.,* no significant perturbations). However, when drought was imposed and integrated with developmental signals, the Propensity distribution plots began to diverge along the x-axis (propensity score) across genotypes. The skewing of propensity plots matched the directional character of fluxes as determined by the positive and negative propensity fractions. Differences in expression fluxes and directional character of the flag leaf transcriptome at booting stage indicated a unique drought response in LPB.

### Candidate yield-associated gene (*OsDEC*) encoded by *qDTY12.1*

Previous study proposed that the major effect of *qDTY12.1* could be explained by a network of genes that regulate root architecture, coordinated by the transcription factor *OsNAM_12.1_* **[23]**. While these findings represent a significant advance in understanding the function of *qDTY12.1-*encoded genes, evidence directly implicating this network to a yield-related mechanism is indirect at best. Guided by the flag leaf transcriptome profiles, we re-examined the expression and annotation of all genes within the *qDTY12.1* syntenic region in the Nipponbare *RefSeq* (chromosome-12) as delineated by the flanking RM28099 and RM511 markers **[20]**.

We found a total of 50 annotated protein-coding genes (**S3 Table)** within the syntenic 1.554 Mbp region in the Nipponbare *RefSeq* within coordinates 15,848,736 bp to 17,401,530 bp **[40]**. However, only 18 of these genes were expressed in at least one developmental stage in any genotype. The expressed genes occurred in small clusters interspersed with genes without any detectable expression in the flag leaf (Fig 4). Co-expression analysis by RiceFREND **[41]** showed that none of the 18 expressed *qDTY12.1* genes formed networks amongst each other, suggesting that none of them were related through a common gene regulon as previously proposed **[23]**. However, the RiceFREND network model showed that 12 of the genes had significant co-expression with other genes from across the genome (Fig 5A).

**Fig 4.**
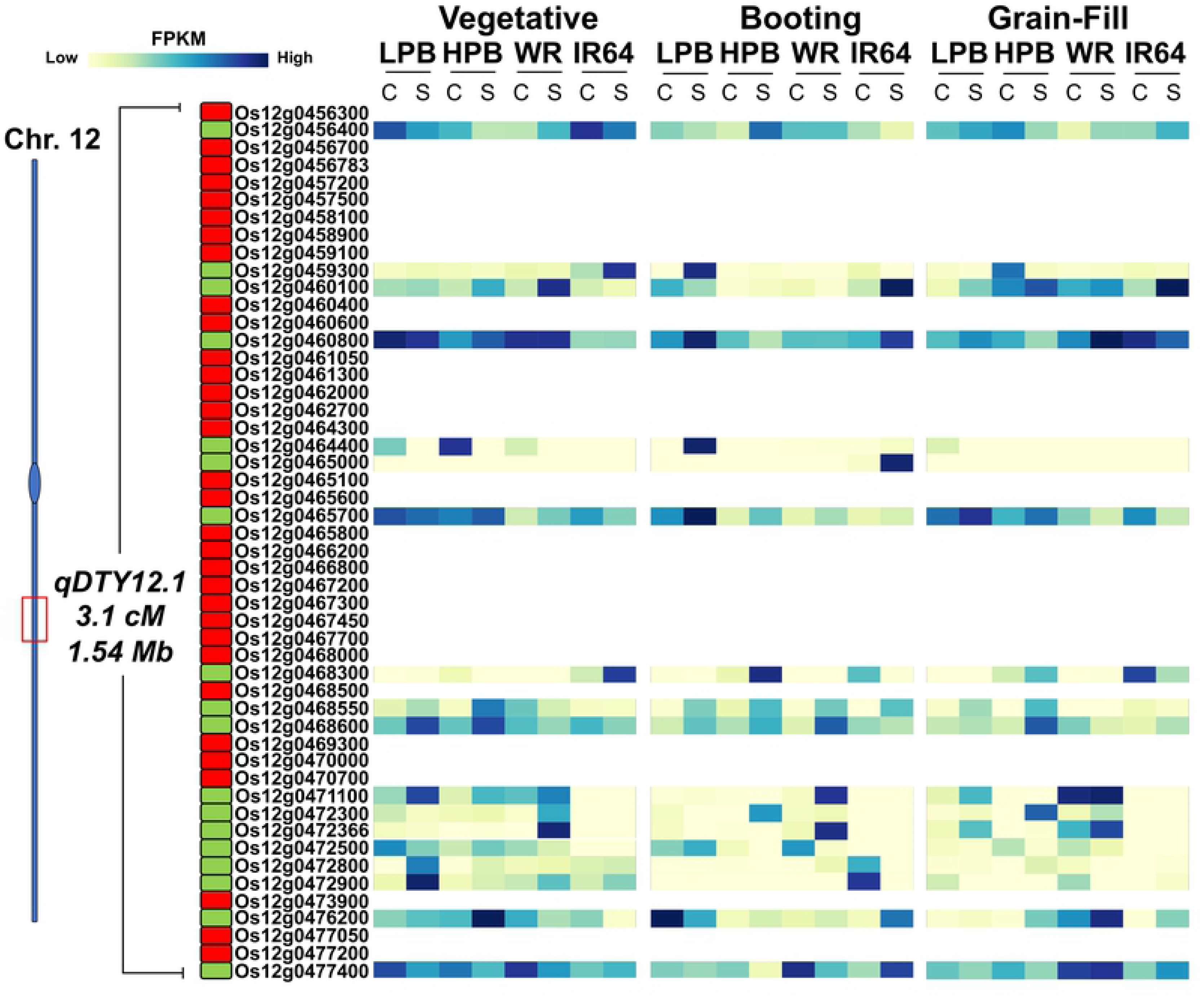
Expression profiles of *qDTY12.1* genes across the spatio-temporal windows of the transcriptomics experiments. Eighteen (18) of the fifty (50) *qDTY12.1* genes had measurable expression. The annotated protein-coding genes were organized by their location on Chromosome-12 (y-axis) and their FPKM-based expression values (yellow = Low; dark blue = High) and plotted across vegetative, booting, grain-filling stages under irrigated and drought conditions (x-axis). Expression is shown for genes with FPKM > 0 (green rectangles) and FPKM = 0 (red rectangles) under irrigated (C – control) or drought (S – stress) conditions.

**Fig 5.**
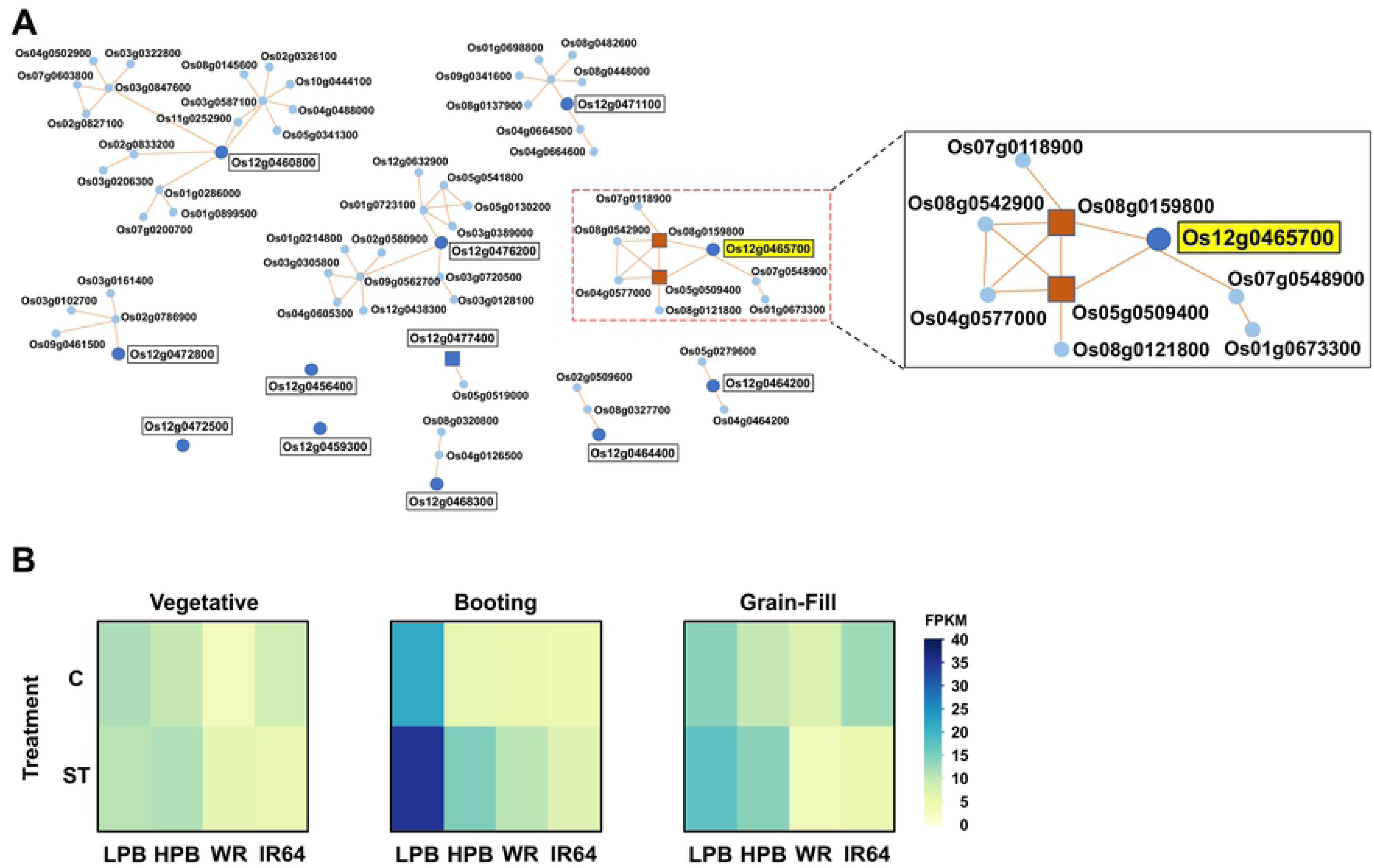
Co-expression of eighteen *qDTY12.1* genes revealed by RiceFREND analysis. (**A**) None of the 18 expressed *qDTY12.1* genes had significant co-expression alliances with each other. However, twelve (12) genes had co-expression alliances with genes outside of *qDTY12.1*, particularly in LPB. The *Os12g0465700* (*OsDEC*) had significant co-expression with two transcription factors (*Os08g0159800, Os05g0509400*) involved in floral meristem functions and singled out as the primary yield-related candidate gene. (**B**) *OsDEC* expression across the genotypic panel at vegetative, booting, and grain-filling stages under irrigated (C) and drought (ST) conditions. *OsDEC* was induced by drought at booting stage only in LPB and first reported to have important roles in cytokinin signaling **[33]**.

One of the genes (*Os12g0465700*) was significantly co-expresssed with two transcription factors (*Os05g0509400*, *Os08g0159800)* whose orthologs in Arabidopsis *(At3g22760* and *At1g32360,* respectively) are involved the regulation of cell division and expansion in floral meristem and expressed mainly in stamens, pollen mother cells, pollen tube, and immature ovules **[42–46]**. The *Os12g0465700* had been previously designated as *DECUSSATE (OsDEC)*, functioning in the regulation of leaf phyllotaxy and associated with the apical meristem (SAM) and root apical meristem (RAM) through cytokinin-mediated signaling **[33]**. It was proposed that *OsDEC* may function as potential transcriptional regulator with broad spectrum targets in response to cytokinin-mediated growth signals **[47–53]**. *OsDEC* was also implicated with reproductive and yield-related functions **[33]**.

*OsDEC* was differentially expressed in the flag leaf under irrigated and drought conditions across developmental stages and genotypes. Differential expression was evident from both the Propensity-based and FPKM-based profiles. *OsDEC* was also significantly upregulated by drought, specifically during the booting stage in LPB but not in HPB, WR and IR64 (Fig 5B). The unique drought-induced expression of *OsDEC* in the flag leaves of LPB at the critical stage of panicle initiation further solidified its potential importance in the regulation of yield-related mechanisms **[54]**.

### Disruption of *DEC* orthologs in Arabidopsis compromised yield

While there is a single copy of *OsDEC* (*Os12g0465700*) in the rice genome, duplicate copies (*At3G03460*, *At5G17510*) have been identified in *Arabidopsis thaliana,* with *At5G17510* as the closest ortholog **(S4 Fig)**. Using the same design of the flowering-stage drought in the rice experiments **(S2 Fig,** Fig 6A), we compared the T-DNA insertion mutants of *At3G03460* (3Gm) and *At5G17510* (5Gm) in Col-0 genetic background with wild-type plants in terms of agronomic performance. Expression of the mutant genes was abolished by T-DNA insertion (Fig 6B).

**Fig 6.**
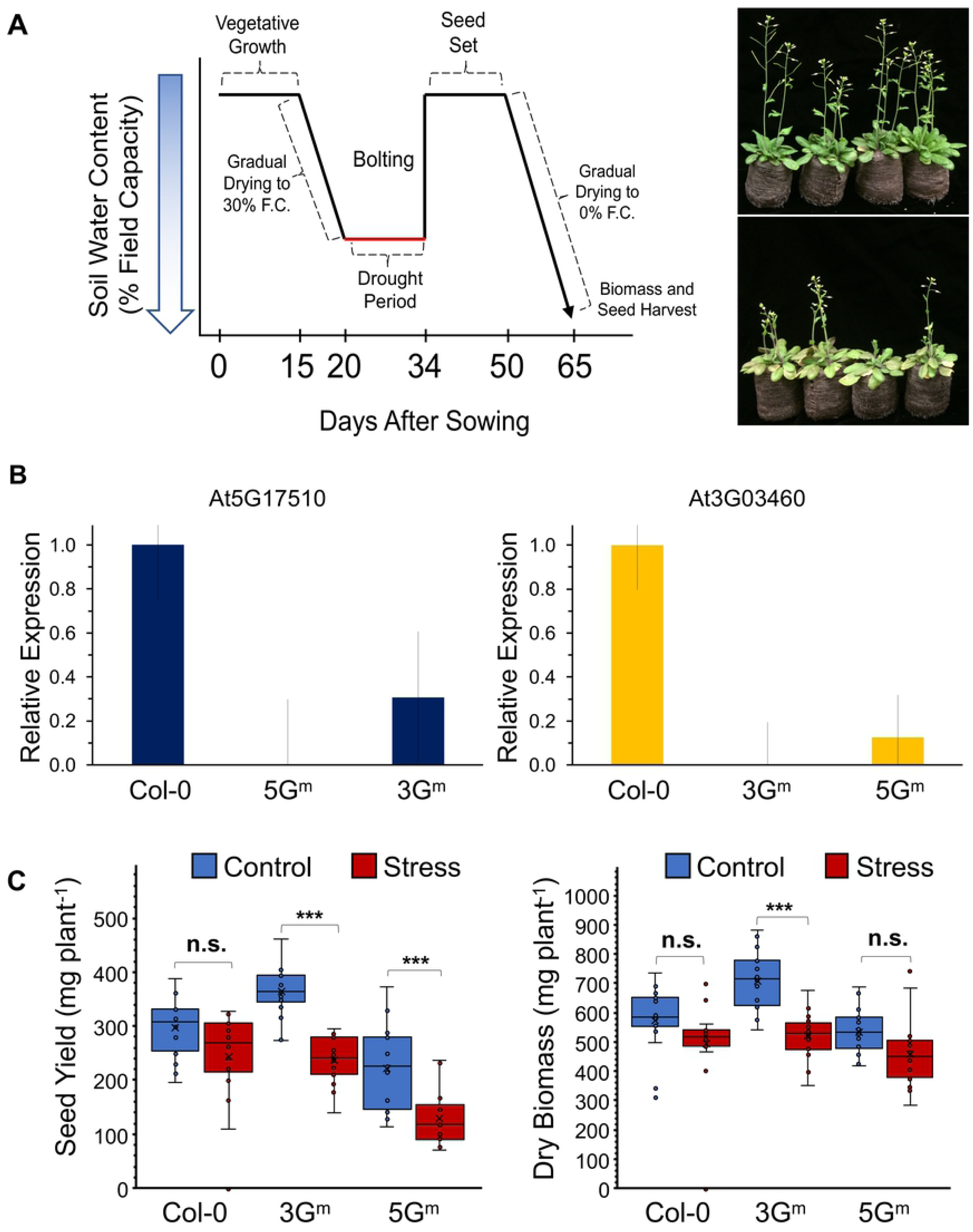
Direct significance of *OsDEC* to yield potential based on heterologous dissection of T-DNA insertion mutants of two orthologous gene copies. (**A**) Growth chamber drought experiments on *Arabidopsis thaliana* ecotype Col-0 and mutants (*At5G17510* = 5Gm, *At3G03460* = 3Gm) mirrored the designs of the drought experiments in rice (S2 Fig). Drought was initiated eight (8) days before bolting (reproductive initiation) and lasted for 14 days, after which plants were re-watered to field capacity until maturity. (**B**) Transcript abundance analysis by qRT-PCR showing the silenced *At5G17510* (5Gm) and *At3G03460* (3Gm) relative to the expression in wild-type Col-0 at day-14. (**C**) Boxplots showing the effects of the loss of *DEC* expression to plant biomass and seed yield. Significant reductions in seed yield under drought are evident in 5Gm and 3Gm (p < 0.001) but not in Col-0, while significant reductions in dry biomass under drought are evident in 3Gm (p < 0.001) but not in Col-0 and 5Gm. Post-hoc comparison of means (all pairs; α = 0.05) was through significant ANOVA using Tukey-HSD: *** significant at p < 0.001.

Results showed that 3Gm was very similar to Col-0 under irrigated conditions in terms of days-to-bolting, days-to-first-bloom, and days-to-seed-set **(S5 Fig)**. On the other hand, 5Gm bolted much earlier and had shorter days to first bloom and seed-set relative to the wild-type Col-0. Strikingly, both 3Gm and 5Gm had significant yield penalties under drought at 34.5% and 41.9%, respectively, while Col-0, having unimpaired drought-induced expression of both *At3G03460* and *At5G17510*, only had 13% yield penalty (Figure 6C**, left panel)**. Analysis of dry biomass showed similar trends as in seed yield, with 3Gm having slightly higher biomass under irrigated condition compared to 5Gm and Col-0 (Fig 6C**, right panel**). However, there was no significant difference in the accumulation of biomass across the three genotypes under drought. The trends in seed yield and dry biomass suggest that differences under drought were perhaps the result of altered source-sink dynamics in the mutants due to the loss of *AtDEC* functions.

### *OsDEC* is the core of a genetic network with other flowering-associated genes

The lack of apparent co-expression of *OsDEC* with other *qDTY12.1* genes suggested that if it was forming a network, the component genes would be located outside of the QTL boundaries. To address this hypothesis, we used *OsDEC* as bait to fish-out for other co-expressed genes in each genotype. In the first step of the iterative procedure, we used the Propensity scores to identify the most significantly co-expressed genes at booting stage, revealing a total of 195 genes in LPB, of which the great majority were cytokinin-related.

In the second step, the primary pool of co-expressed genes was further reduced to a much tighter cluster of 30 genes with the common gene ontology (GO) keywords of ‘*cytokinin’*, ‘*flowering’*, and ‘*inflorescence*’ (Figure 7A**, red box)**. With a threshold Propensity value of n ≥ 0.5, the core of the network with the most significant similarities in flux with *OsDEC* was identified as a smaller subset of 11 genes (**Table 1**). The FPKM-based co-expression profiles of this core is shown in the hierarchical clustering in Fig 7B, with Clades-2, −3, and −4 exhibiting the most highly significant co-expression with *OsDEC* under both irrigated and drought conditions. *OsDEC* is a member of Clade-2 with three other genes (*Os07g0108900 = OsMADS15*; *Os05g0521300 = OsPHP3*; *Os03g0109300 = OsLOGL3*). Annotation queries indicate that these genes shared common functions by virtue of their roles in the specification of ‘*inflorescence meristem identity*’ (GO:0048510), ‘*floral organ regulation*’ (GO:0048833), ‘*cytokinin signaling*’ (GO:0009736), and ‘*cytokinin biosynthesis*’ (GO:0009691). The only gene in Clade-3 was annotated as a floral organ regulator (Os07g0568700; GO:0048833). The other solitary gene in Clade-4 is annotated as *OsMADS-14* (*Os03g0752800*), which functions in the specification of inflorescence meristem identity (GO:0048510).

**Fig 7.**
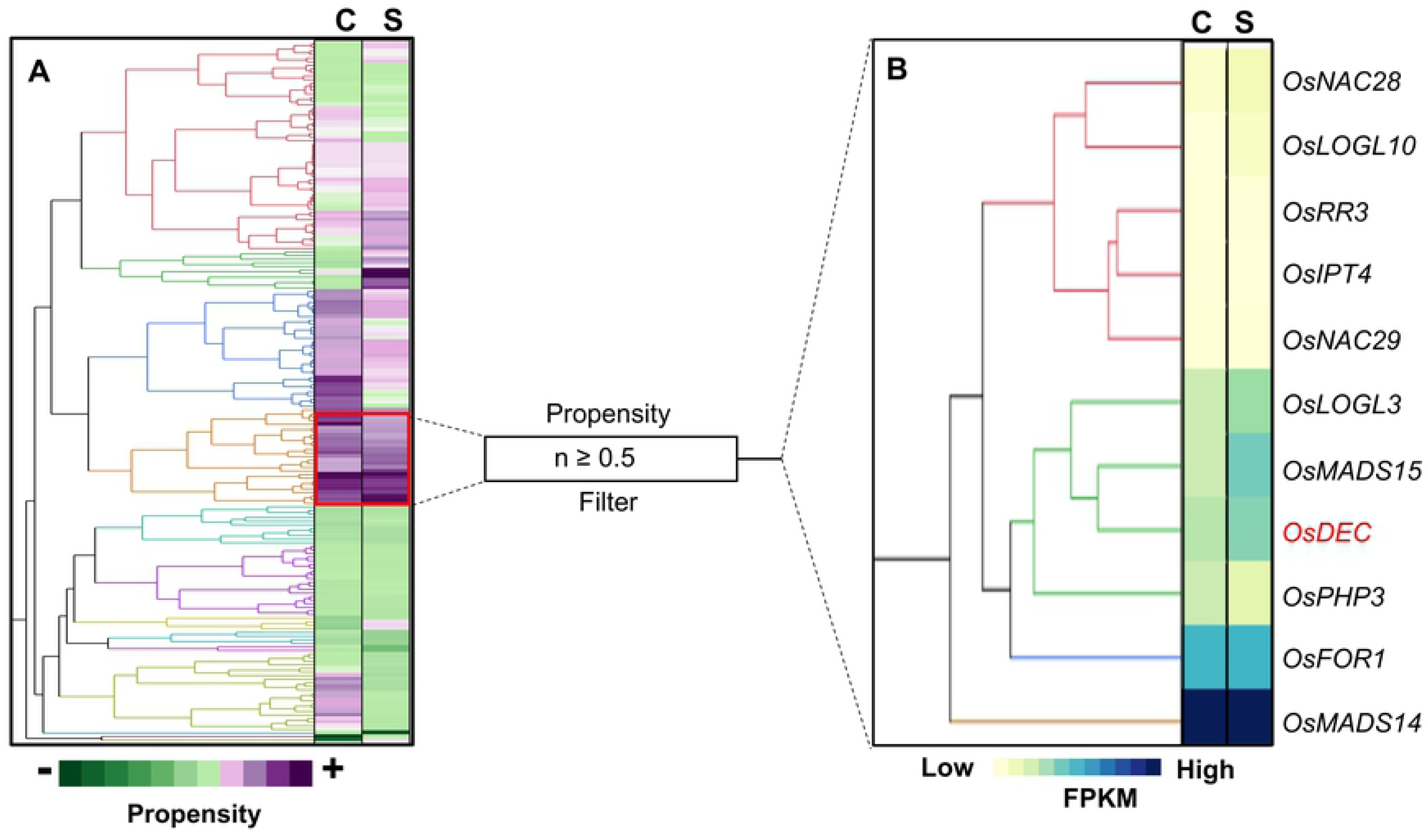
Co-expression of *OsDEC* with genes involved in the regulation of flowering at booting stage in LPB. (**A**) Hierarchical clustering of Propensity values for *OsDEC* and other genes (n = 195) associated with cytokinin signaling. A robust cohort of 30 genes with common ontology (GO) of cytokinin, flowering, and inflorescence were extracted (red box) from the LPB transcriptome. Refinement using a Propensity threshold of n ≥ 0.5 identified 11 genes with highly significant co-expression. (**B**) Hierarchical clustering of FPKM-based expression established the six-gene network hub, comprised of *OsDEC*, *OsLOGL3*, *OsMADS15*, *OsPHP3*, *OsFOR1*, and *OsMADS14* characterized by GO for cytokinin signaling, inflorescence meristem identity, and floral organ regulation.

**Table 1.** List of genes included in the main hub of the DEC-network and used as baits for extracting the components of the full DEC-network at booting stage.

| **Locus ID** | **\*Oryabase Gene Symbol** | **†RAP-DB Description** |
| --- | --- | --- |
| ***Os02g0555300*** | *OsNAC28* | No apical meristem (NAM) protein domain containing protein. |
| ***Os02g0830200*** | *OsRR3* | A-type response regulator, Cytokinin signaling |
| ***Os03g0109300*** | *LOGL3* | Similar to Lysine decarboxylase-like protein |
| ***Os03g0752800*** | *OsMADS14* | Similar to Isoform 2 of MADS-box transcription factor 14. APETALA1 (AP1)/ FRUITFULL (FUL)-like MADS box transcription factor, Specification of inflorescence meristem identity |
| ***Os03g0810100*** | *OsIPT4* | Similar to tRNA isopentenyl transferase-like protein (Adenylate isopentenyltransferase) |
| ***Os05g0521300*** | *OsPHP3* | Similar to Histidine-containing phosphotransfer protein 4 |
| ***Os07g0108900*** | *OsMADS15* | Similar to MADS-box transcription factor 15. APETALA1 (AP1)/ FRUITFULL (FUL)-like MADS box transcription factor, Specification of inflorescence meristem identity, sexual reproduction |
| ***Os07g0568700*** | *OsFOR1* | Polygalacturonase-inhibiting protein, Inhibitor of fungal polygalacturonase, Regulation of floral organ number |
| ***Os08g0115800*** | *ONAC29* | NAC transcription factor, Regulation of cellulose synthesis |
| ***Os10g0479500*** | *LOGL10* | Similar to carboxy-lyase |
| ***Os12g0465700*** | *DEC* | Plant-specific protein containing a glutamine-rich region and a conserved motif, Controls of phyllotaxy by affecting cytokinin signaling |

†RAP-DB – The Rice Annotation Project Database (https://rapdb.dna.affrc.go.jp/index.html)
\*Oryabase – Integrated Rice Science Database (https://shigen.nig.ac.jp/rice/oryabase/)

To capture the secondary and tertiary components of the *DEC*-network, other genes with cytokinin-associated functions that exhibited significant co-expression with *OsDEC* and/or its five (5) other direct cohort genes were identified in the third iteration. FPKM-based hierarchical clustering revealed a larger group that formed thirteen clades of tightly co-expressed genes around the *DEC*-network. A total of 36 genes (**S4 Table**) that were most significantly co-expressed with *OsDEC* and its direct cohort genes were contained within two clades that reflect the potential functional significance of the *DEC*-network (Fig 8A). The main hub of this network of 36 genes is *OsDEC* itself and two MADS-box transcription factors that regulate meristem transition from vegetative to flowering stage, *i.e., Os07g0108900* = *OsMADS15*, *Os01g0922800* = *OsMADS51* (Fig 8B). The other ‘*peripheral*’ components surrounding the *DEC*-network were dispersed across seven clades, all of which are associated with vegetative to reproductive transition of the meristem.

**Fig 8.**
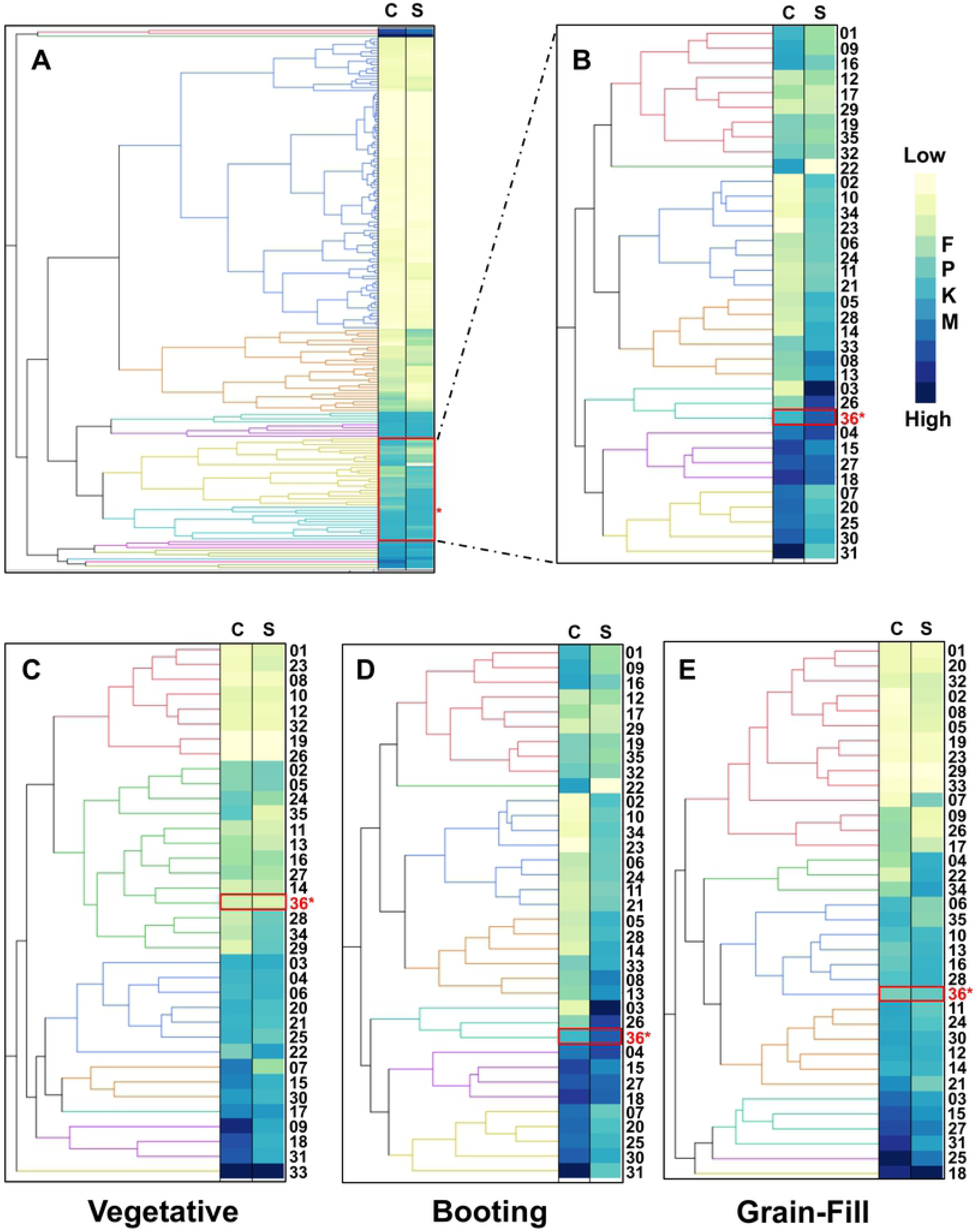
Differential organization of *DEC*-network showing the uniqueness of LPB. High similarities in *OsDEC* network were evident across all genotypes at vegetative and grain-filling stages. (**A**) Hierarchical clustering of FPKM-based transcript abundances revealed 13 clades of co-expressed genes surrounding the *DEC*-network hub. Clades-7 and −8 (red box with asterisk) contained 36 genes that were highly co-expressed with *OsDEC*. (**B**) Final composition of the *DEC*-network of LPB based on hierarchical clustering of FPKM-based transcript abundances. The ‘core’ of the network consisted of *OsDEC* (36*), *OsMADS15* (26), and*OsMADS51* (03), all of which are directly involved with meristem transition from vegetative to reproductive. The other 33 genes formed the peripheral components with direct linkages to reproductive functions. (**C-E**) Hierarchical clustering of FPKM-based transcript abundances across the 36-member *DEC-*network. Numbers to the right of dendograms (Locus ID, position, annotation, etc.) are detailed in S4 Table. Red asterisk marks the position of *OsDEC*. C – Control/irrigated; S – Stress/drought.

### *DEC*-network is specific to booting stage and genotype-dependent

To further understand the significance of *OsDEC* to yield maintenance under drought, we compared the *DEC*-network organization across developmental stages within LPB, *i.e.,* vegetative versus booting versus grain-filling, and across genotypes with or without the *qDTY12.1*, *i.e.,* LPB versus HPB, donor parent WR, and recipient parent IR64. Hierarchical clustering showed significant differences in co-expression among the 36 ‘*core*’ and ‘*peripheral*’ genes that comprise the *DEC*-network across developmental stages (Fig 8 **C-E)**. In LPB, genes of the *DEC*-network were coordinately induced by drought specifically at booting stage, while no significant changes in expression were detected at vegetative and grain-filling stages.

Further examination of the organization of the booting-stage network across genotypes revealed widely divergent patterns, with only LPB showing evidence of coordinated expression of all *‘core’* and *‘peripheral’* components (Fig 9A). The genotype-dependent and booting stage-specific signatures in LPB suggested that the operability of the *DEC*-network was likely a consequence of the proper alignment of all the upstream regulatory components that established the optimal expression of *OsDEC* and subsequently all of its downstream cohort/peripheral genes.

**Fig 9.**
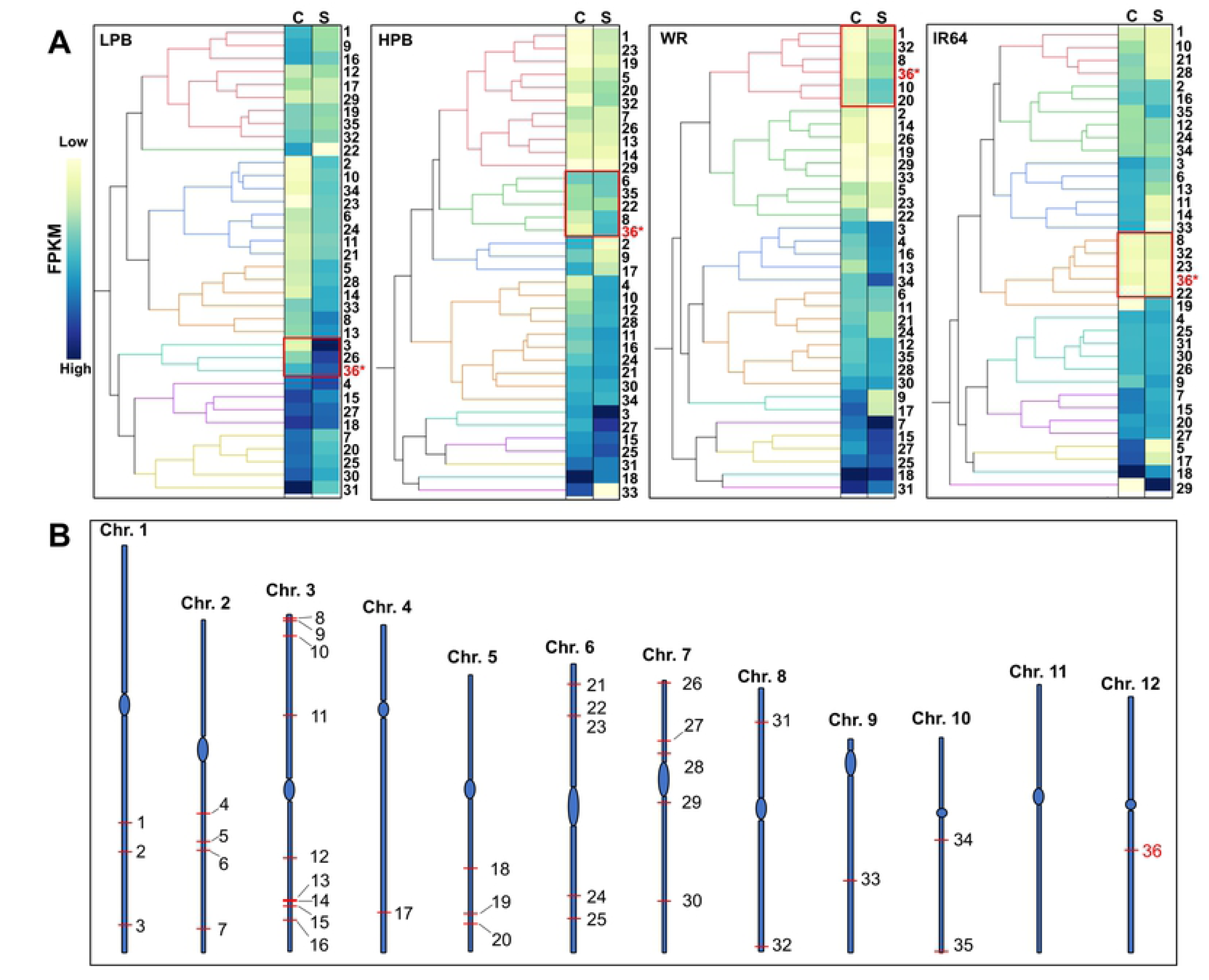
Organization of the booting stage *DEC-*network across genotypes. The *OsDEC* formed networks with other genes in the genetic background, and the network is highly organized in LPB but not in the other genotypes, where homologous networks appeared fragmented and disorganized. (**A**) The organization and expression character of the *DEC-*network at booting stage are distinct in each genotype. In LPB, the network is characterized by an inductive pattern while a static pattern was evident in HPB, WR, and IR64. (**B**) Distribution of the members of the functional *DEC*-network across the rice genome outside of *qDTY12.1*. Numbers to the right of the dendograms are described (Locus ID, position, annotation, etc.) in S4 Table. Number with red asterisk indicate the position of *OsDEC*. C – Control/irrigated; S – Stress/drought.

The disorganized *DEC*-network in HPB appeared to suggest the opposite of what was observed in LPB, perhaps due to the lack of complementary alleles for the upstream components that facilitate the same level of network organization as observed in LPB. The variant patterns in the *DEC*-network between the sibling LPB and HPB, both of which had the same *OsDEC* allele from WR **(sequence data not shown)**, further implied an efficient integration of stress and developmental signals through the interaction between *qDTY12.1* and its *‘peripheral’* cohort genes in the genetic background (Fig 9B). Thus, a complete *DEC*-network appeared to be strategic to an optimal integration of drought-mediated signals with developmental signals during the early stages of flowering when the critical reproductive sink is being established.

### Yield component traits associate with the *DEC*-network

For further interpretation of the larger biological significance of the *qDTY12.1*-encoded *OsDEC* and its network with other genes in the genetic background, we established a biological network map through the Knetminer knowledge integration tools **[55]**. This analysis links many pieces of relevant information from all types of genetic studies curated in the literature to establish direct or indirect associations between a gene or network of genes and physiological and agronomic traits. The knowledge integration map directly linked all but one of the 36 genes that comprised the *DEC*-network with various yield component traits, particularly those relevant to source-sink regulation, sucrose and starch biosynthesis and deposition, grain-filling, seed development and maturation, and seed weight (Fig 10**, S5 Table)**.

**Fig 10.**
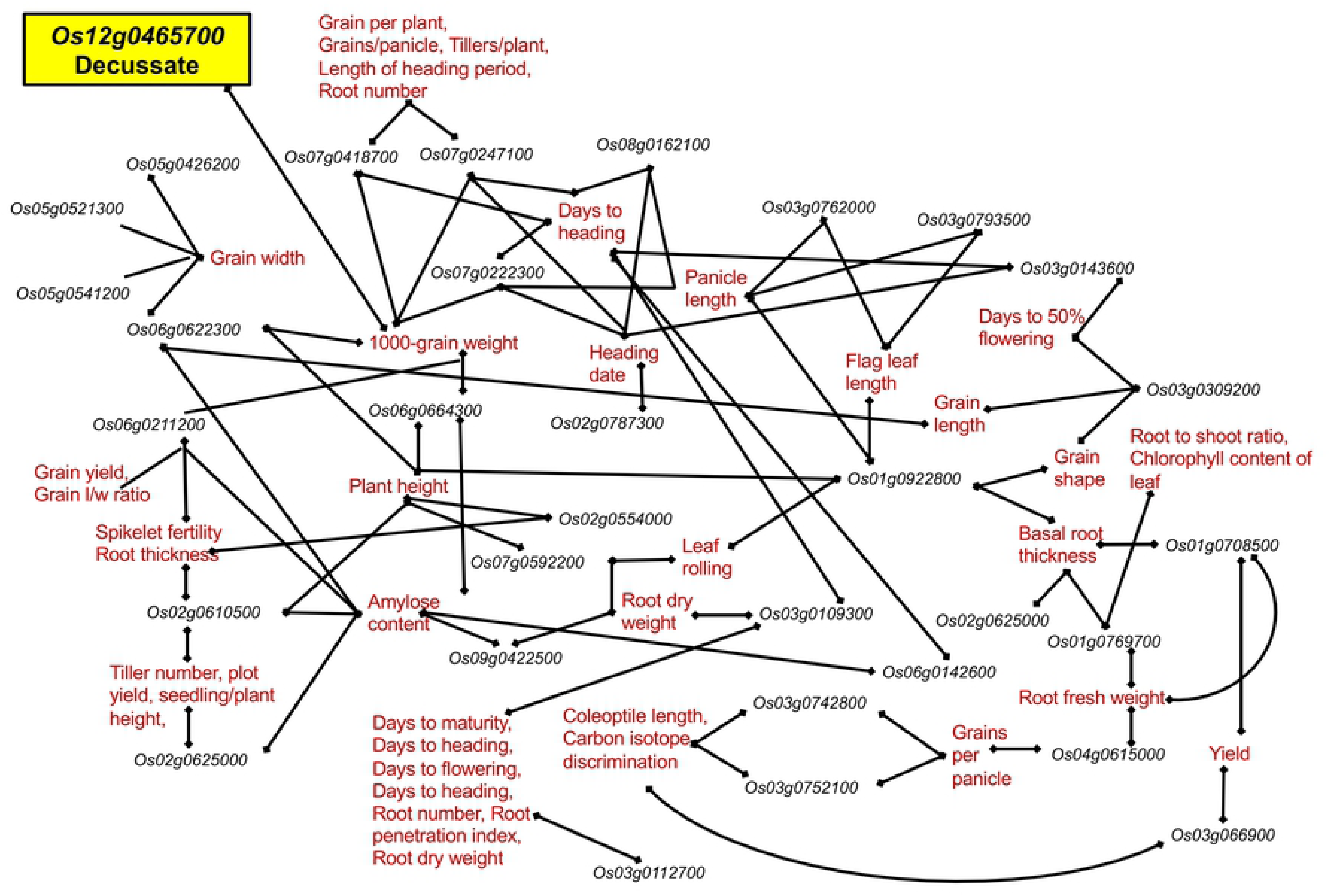
KnetMiner knowledge integration map depicting the biological functions associated with the operative booting stage-specific *DEC*-network in LPB. The knowledge integration map linked all but one of the 36 genes to various yield component traits including grains per plant, tillers per plant, grain length and width, panicle length, amylose content, carbon isotope discrimination, days to flowering, days to heading, spikelet number and fertility, and grain yield. The complete list of traits generated by the KnetMiner are summarized in S5 Table.

A recent study in maize highlighted the significant impacts of *ZMM28* overexpression to flowering time, plant growth, photosynthetic capacity, nitrogen utilization, and yield under drought **[56]**. *ZMM28* is a member of the AP1-FUL sub-group of MADS-box transcription factors with critical roles in the regulation of flowering time, floral organ identity, and vegetative to reproductive transition **[57–59]**. We found that *OsMADS18* (*Os07g0605200*), the closest ortholog of *ZMM28* in rice, along with two other MADS-box genes (*Os03g0752800* = *OsMADS14; Os07g0108900* = *OsMADS15*) had strikingly similar expression as *OsDEC* in LPB at the booting stage (Fig 11). Expression peaked at booting stage in LPB and IR64, but not in HPB and WR. These findings suggested the influence of IR64 genetic background in the optimal configuration of *DEC-*network in LPB but not in HPB. Of important note, the expression of *OsMADS18* in LPB across developmental stages was very similar to the *zmm28* signature in transgenic maize **[56]**.

**Fig 11.**
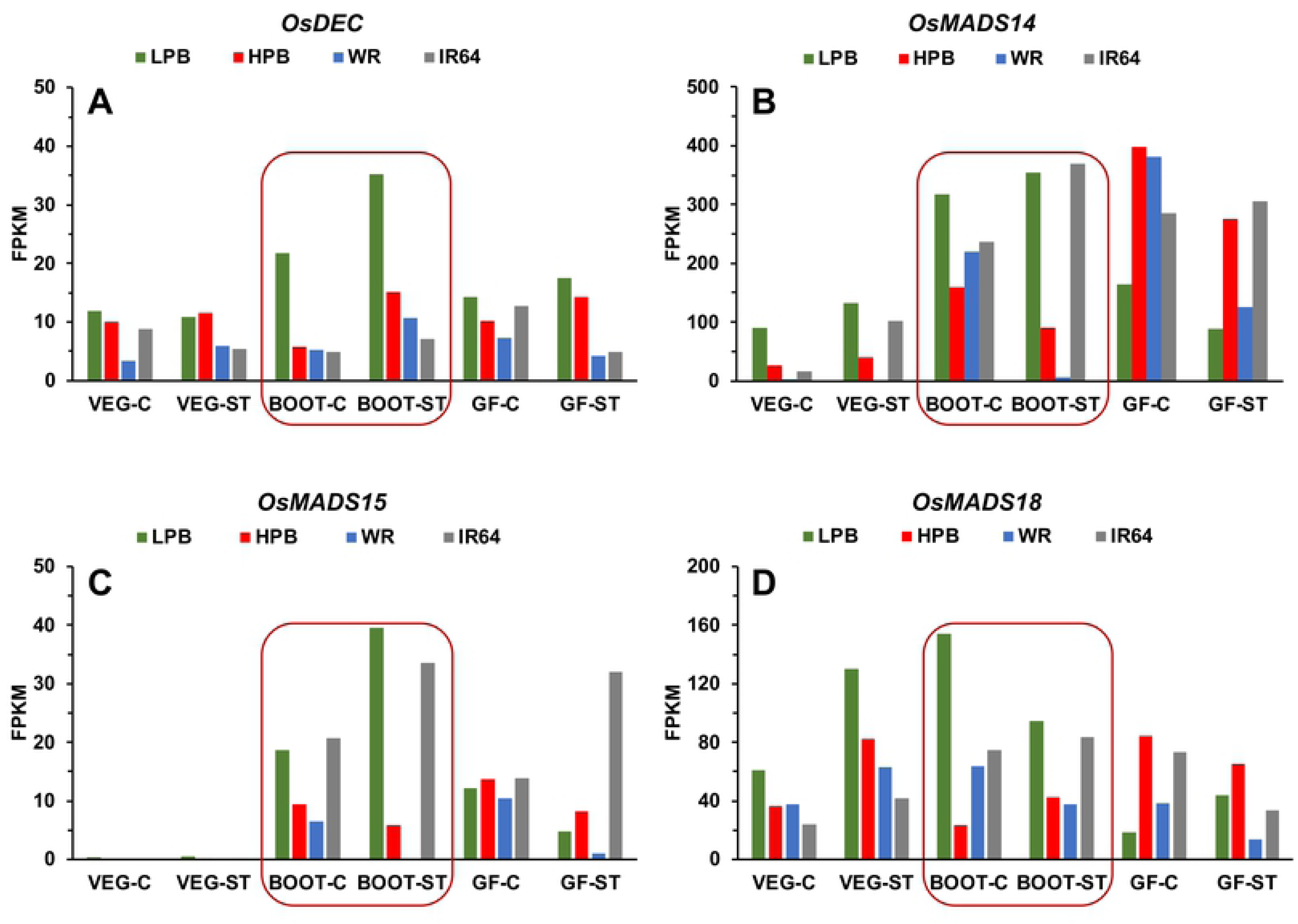
Expression of critical MADS-box transcription factors at booting stage in LPB mimic the signature of *OsDEC*. The temporal expression of *OsDEC* and three MADS-box transcription factors point to a mechanism for regulating flowering-time under drought. (**A-D**) FPKM-based expression plots of *OsDEC*, *OsMADS14*, *OsMADS15*, and *OsMADS18* across growth conditions (irrigated, drought) and developmental stages. *OsMADS14*, *OsMADS15*, *OsMADS18* (documented to be intimately involved in flowering and meristem identity) were induced by drought at booting stage (red boxes) in LPB and IR64, but not in HPB and WR. Expression of *OsMADS18* across developmental stages mimicked the overexpression (OE) of *ZMM28* (maize ortholog) that led to improved growth and yield **[56]**. VEG-C (vegetative control/irrigated); VEG-ST (vegetative stress/drought); BOOT-C (booting control/irrigated); BOOT-ST (booting stress/drought); GF-C (grain-filling control/irrigated); GF-ST (grain-filling stress/drought).

LPB had the shortest delay, *i.e.,* eight (8) days, in flowering time under drought in comparison to IR64, HPB, and WR, with ten (10), sixteen (16), and eighteen (18) days delay, respectively (Fig 1). Coupled with the observed trends in MADS-box expression, it appeared that the *DEC*-network in LPB had integrated the function of *OsMADS14, OsMADS15,* and *OsMADS18* towards a mechanism for reducing time delays in reproductive growth transition during drought.

## Discussion

Introgression and pyramiding of large-effect QTLs such as *qDTY12.1* have shown major incremental improvements in rice yield maintenance under drought **[14,15,17,19,28,60–62]**. However, there have been instances when the introgression of *qDTYs* did not confer the expected phenotypic effects **[22]**. A similar phenomenon has been reported with the introgression of the *SalTol* QTL for salinity tolerance in different rice cultivars, when the presence of the QTL alone did not necessarily lead to the expected phenotypic effects **[30,63]**. Inconsistent effects are caused by negative or positive epistatic interactions between the QTL genes and other genes in the genetic background that could either enhance or drag the effects of the QTL **[7,8]**. In this study, we illuminated this enigma by integrating the new concepts of the *Omnigenic Theory* **[31]**, and by using the mechanisms that cause transgressive traits in rice to further illuminate our conceptual framework **[7,8,30]**.

### Significance of *qDTY12.1* to genetic network rewiring

It was postulated that non-parental traits created by genetic recombination are due to genetic coupling-uncoupling and network rewiring effects. Rewired genetic networks are caused by large assemblages of synergistic or antagonistic genes that get coupled or uncoupled during multiple rounds of recombination. In the context of the *Omnigenic Theory*, the few ‘*core’* genes with major effects on phenotypic variance could either be coupled or uncoupled with numerous compatible or incompatible *periphera*l genes with minute but additive effects on the phenotypic variance **[31]**. The additive effects of the ‘*peripheral’* genes across the genetic background may either enhance or drag the effects of the ‘*core’* genes that function as the hub of the network.

Our results showed yet another layer of evidence that the inconsistent effects of *qDTY12.1* observed across two sibling introgression lines in the genetic background of IR64 were due to either optimally (LPB) or sub-optimally (HPB) rewired genetic networks, with a *qDTY12.1*-encoded regulatory gene *OsDEC* functioning as the hub of the network. We hypothesized that while backcross introgression of the functional *qDTY12.1* allele from Way Rarem (WR) into IR64 genetic background (through a bridge donor derived from WR x Vandana) may have preserved the integrity of the original *qDTY12.1* allele by marker-assisted selection of the foreground, the genomic environments (background) of the introgressed *qDTY12.1* were likely to be significantly divergent between sibling introgression lines. By extension, the rewired genetic networks were configured by many loci/alleles from either parents, organized in such a manner that either optimal and sub-optimal alliances define the operative structure of the network. Further, the superior progeny (LPB) appeared to contain not only the required network hub, that is the *OsDEC* allele from WR, but also the optimal assemblage of ‘*peripheral’* alleles across the genetic background leading to a fully functional synergy. These ‘*peripheral’* alleles are likely to have come directly either from IR64 or remnant and cryptic introgression of alleles from WR or Vandana that escaped the resolution and scope of marker-assisted selection of the background genome. On the other hand, while the inferior sibling HPB also contained the identical network core from WR (*OsDEC*), it appeared to be lacking the same optimal assemblage of ‘*peripheral’* alleles from across the genetic background to configure a functioning synergy for the *DEC*-network (**S1 Fig)**.

Comparative dissection of the flag leaf transcriptomes of LPB and HPB in relation to the *qDTY12.1* donor parent WR and recipient parent IR64 showed that while the global patterns under irrigated condition at the vegetative and grain-filling stages were generally similar across the genotypes, there were drastic differences at the booting stage (Fig 2). These differences appeared to be the results of coupling-uncoupling effects, hence interaction of distinct subsets of synergistic and antagonistic alleles from either parent. As such, the positive effect of *qDTY12.1* introgression in context of *DEC*-network would be manifested only when optimal number of compatible ‘*peripheral’* alleles with additive effects are assembled to generate the transgressive genetic network that was apparent in the booting-stage transcriptome of LPB. It is evident based on the distinct transcriptomic signatures of LPB and HPB, that while *qDTY12.1* has a large effect on yield, expressing its full potential requires many other ‘*peripheral’* genes across the genetic background.

Our current results do not indicate that any other genes within the *qDTY12.1* are important for the full functionality of the *DEC*-network. Indeed, all of the 35 peripheral genes that comprised the functional *DEC*-network in LPB are dispersed throughout the genome, clearly outside of the boundaries of *qDTY12.1* (Fig 9B). Thus, the transgressive nature of yield maintenance under drought as conferred by *OsDEC*, requires a synergy with many other genes in the genetic background. This was made abundantly clear by the fact that although HPB and WR had the same *qDTY12.1* allele as LPB, their yield potentials under drought were woefully inferior.

Another important advance contributed by this study is the discovery that while LPB and HPB were assumed to be largely similar with regard to *qDTY12.1*, the flag leaf transcriptome of LPB specifically at booting stage was drastically different from its recurrent and QTL donor parents and sibling introgeression line (Fig 2B). Booting stage represents a critical crossroad of photosynthetic source-sink dynamics between the flag leaf and developing inflorescence, characterized by physiological and biochemical processes that sustain seed development **[64–68] (S2 Fig)**. As such, events unique to LPB at booting stage provides a valuable link to the functional significance of *qDTY12.1* to cellular mechanisms critical to yield components. It has been shown that the timing of drought at the initiation of booting is most deleterious, with negative effects on yield-related traits including grain number per panicle, panicles per area, and total above ground biomass **[39]**. The significance of *qDTY12.1* is consistent with the synchronized activation of the *DEC*-network when drought coincides with the early stages of floral organ development **[17,18,25,69]**.

### *OsDEC* affects yield-related processes through the cytokinin signaling pathway

The *OsDEC* was singled out as the most likely candidate for a yield-related gene of *qDTY12.1* based on its unique drought-induced expression in the flag leaf of LPB but not in the other genotypes, specifically at the initiation of booting. We found that *OsDEC* was the only one among the 18 *qDTY12.1* genes transcribed in the flag leaf that was also differentially induced by drought at the booting stage only in LPB with significant co-expression with two transcription factors related to reproductive growth (Fig 4, Fig 5 **A-B)**. While the specific biochemical function of *OsDEC* remains unknown, it is known to have a regulatory function over Type-A and Type-B Response Regulators (ARR) in the two-component cytokinin signal transduction pathway **[33,70–72]**. Cytokinin is intrinsic to a myriad of cellular processes that are critical for seed development as well as for mediating cellular signals in response to drought **[47,49–52,73,74]**. Studies in many agronomically important crops have also shown that overexpression of cytokinin biosynthetic genes leads to significant improvements in yield potential under drought **[53,75–77]**.

Results of this study support a hypothesis that through a cytokinin-mediated pathway, *OsDEC* regulates physiological processes in the flag leaf that appeared to be important in adjusting the timing of floral organ initiation when the photosynthetic source is perturbed by drought. We further infer that this mechanism could be important in ensuring the early establishment of a strong reproductive sink to sustain the requirements of seed development and maturation when resources continue to be limited by drought effects. Indeed, the 35 other genes in the *DEC*-network were mostly regulatory genes with key functions in the regulation of floral meristem, vegetative to reproductive transition, cytokinin signal transduction, and other aspects of reproductive growth. These trends were further reiterated by the models generated by KnetMiner, which showed that all genes in the larger *DEC*-network funnel into processes involved with seed development, grain filling, sucrose transport, starch biosynthesis, and many other yield-component traits (Fig 10).

Furthermore, many introgression lines of *qDTY12.1* have been extensively studied to determine what physiological characteristics are important in the maintenance of low-yield-penalty under drought **[25–27,78,79]**. These characteristics include water uptake efficiency, increased proline levels in roots, improved remobilization of amino acids for nitrogen status, improved transpiration efficiency, increased panicle branching, increased lateral root formation, and a reduction in flowering delay under drought. These characteristics are consistent with the central role of *OsDEC* in integrating survival, developmental and stress-related responses to minimize the cost of cellular perturbations to reproductive growth **[22]**.

From the standpoint of productivity, flowering represents a developmental crossroad. As such, it is regulated tightly by environmental signals to ensure reproductive success of the species, hence the process is dynamic, multi-faceted, and with multiple levels of control over a large number of genes. A closer examination of the components of the functional *DEC*-network (*i.e.,* 35 genes) indicate direct connections to one or more molecular, cellular, or biological functions that are relevant to the control of flowering time, including *hormonal signaling* (GO:0007267), *light signaling* (GO:0009416), *epigenetic control* (GO:0040029), *developmental control of floral organ differentiation and fate* (GO:0048437), *maintenance of reproductive meristems* (GO:0010073), and *transcriptional regulation* (GO:0006357). Some of the well-known MADS-box transcription factors such as *OsMADS14*, *OsMADS15*, and *OsMADS18* define the hallmark signatures of direct association of *OsDEC* with the regulation of flowering time **[59,80,81]**.

The magnitude of drought-induced delay in flowering is strongly correlated with yield retention in rice **[82,83]**. Progressive drought imposed before the onset of flowering caused 8, 16, 18, and 10 day delays in normal flowering time in LPB, HPB, WR, and IR64, respectively (Fig 1). Under limited water conditions, earlier flowering would provide a developmental adjustment to minimize the effects of continuous depletion of photosynthetic sources that would normally sustain reproductive transition. Therefore, expression of many flowering-related genes with molecular and cellular functions associated with *floral organ identity* (GO:0010093), *inflorescence meristem maintenance* (GO:0010077), and *spikelet development* (GO:0009909) appeared to commence earlier in LPB due to drought. These GO terms are relevant to the establishment and maintenance of critical yield-component traits such as *number of fertile spikelets*, *number of reproductive tillers*, *number of panicles*, *grain weight*, *number of grains per panicle*, and *panicle size,* as shown in the KnetMiner Map and verified by yield components data (Fig 10, Fig 1).

### Potential implications of the *DEC*-network at the molecular and cellular levels

In earlier efforts to characterize the cellular functions of *OsDEC* using *dec* mutants, the following conclusions emerged: 1) *OsDEC* is insensitive to exogenous cytokinin; 2) *OsCKX2* and other *cytokinin oxidase* genes were upregulated in knock-out mutants; 3) active cytokinins cZ and iP, along with some of their intermediates were significantly reduced in mutants; 4) expression of *LOG* (*Lonely GUY*) genes were not affected in mutants; and 5) Type-A Response Regulators were downregulated while some Type-B Response Regulators were upregulated **[33]**. Additionally, *OsDEC* is most highly expressed in immature leaves and inflorescence apex. DEC protein potentially functions as transcriptional regulator based on the N-terminus glutamine-rich domain associated with chromatin remodeling functions **[84– 88]**. By integrating these information with other co-expressed genes in the flag leaf transcriptome of LPB, we propose a hypothetical model of the mechanisms by which the *DEC*-network regulates early flowering (Fig 12).

**Fig 12.**
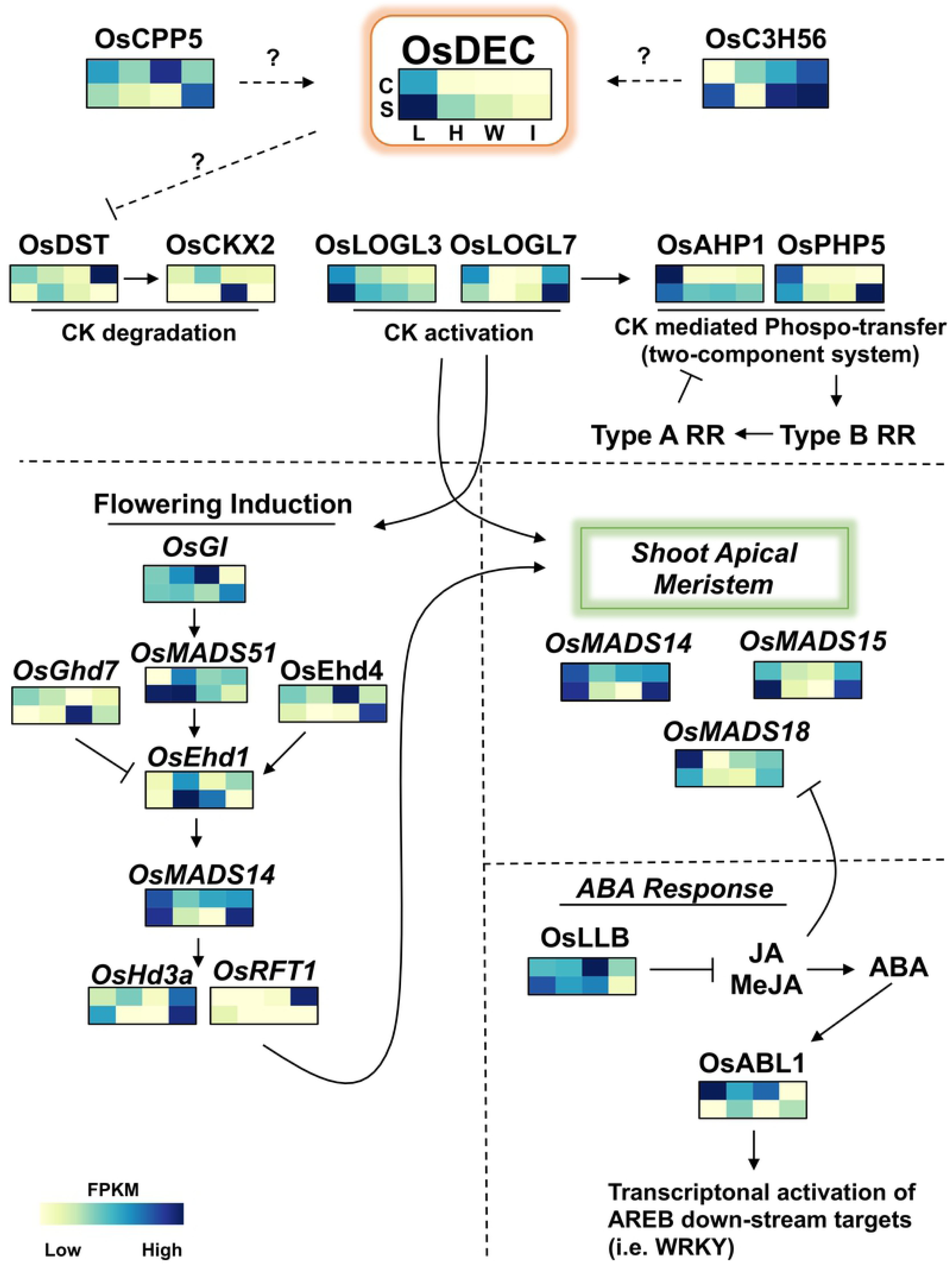
Putative molecular mechanism of the *DEC*-network modeled through the integration of relevant information from the literature with the trends uncovered from the flag leaf drought transcriptomes. In concert with other genes (peripheral) across the genetic background, *OsDEC* anchors a network that effectively mediates early transition to reproduction, thereby facilitating processes critical for grain productivity under drought. Transition of the meristem from vegetative to reproductive stage is mediated by cytokinin through the phospho-transfer system (*OsAHP1, OsPHP5*) leading to the enhancement of active cytokinin pools through the expression of the biosynthetic genes *OsLOGL3* and *OsLOGL7*, and concomitant suppression of *OsCKX2* involved in degradation. Induction of spikelet development is promoted by *OsMADS14*, *OsMADS15*, *OsMADS18*, and *OsMADS51*, and *OsHd3a* (florigen), which trigger the early onset of flowering under drought. Early formation of reproductive sink efficiently redirects the photosynthate to reproductive processes. The ‘*taming*’ effect of ABA response (*OsLLB*, *OsABL1*) prevents unnecessary wastage of photosynthates that leads to large trade-offs to yield. C – control/irrigated; S – stress/drought; L – LPB; H – HPB; W – WR; I – IR64.

We hypothesize that in LPB, the pools of active cytokinins would be enhanced due to drought-mediated upregulation of *OsLOGL3* and *OsLOGL7* (*Lonely-Guy*) and downregulation of *OsCKX2*. The significance of *OsCKX2* downregulation to the enhancement of grain yield in rice has been confirmed **[73]**. In the model, the pool of active cytokinins is upregulated with concomitant downregulation of cytokinin degradation by *OsCKX2*. Studies have shown that Os*DEC* regulates *CKX* expression but not *LOG* expression **[33]**. It has also been reported that a drought and salinity-associated C2H2 zinc-finger transcription factor (*OsDST*) is directly involved in the regulation of *OsCKX2* **[89]**. It has also been reported that *DST* mutation (*OsDST^reg1^*) downregulates *OsCKX2*, thereby cinreasing the level of active cytokinin **[90]**. Downregulation of *OsDST* was evident in the flag leaf transcriptome at the booting stage, with −2.2 and −5.9 log2-fold decreases in transcript abundance under drought in LPB and IR64, respectively. In contrast, *OsDST* was upregulated in HPB and WR with 0.74 and 0.62 log2-fold changes, respectively **(S6 Fig)**.

Increased levels of active cytokinin have been implicated to yield enhancement in rice, which correlates well with the higher yield potential of LPB under drought and parallel upregulation of cytokinin biosynthetic genes and downregulation of cytokinin degradation genes such as *OsCKX2* **[49,50,52]**. Additionally, cytokinin signaling directly affects other genes that regulate flowering **[48,91–93]**. Accumulation of active cytokinin in LPB suggests a mechanism that facilitates earlier induction of flowering under drought as a penalty-avoidance response by establishing proper source-sink dynamics earlier before the source becomes more limited or depleted.

Network of *OsDEC* with *OsMADS14*, *OsMADS15*, and *OsMADS51* showed that indeed the flowering pathway was induced earlier in LPB. These MADS transcription factors are critical for regulating inflorescence meristematic processes in rice **[59,80,94,95]**. A recent study in maize also showed that overexpression of the *OsMADS18* ortholog in maize (*zmm28)* led to significant increases in yield under sub-optimal irrigation **[56]**. It has also been shown that *OsMADS18* accelerates the transition of meristem from vegetative to reproductive by promoting the florigen *Hd3a* via cytokinin signaling **[81,96]**. It has been reported that methyl-jasmonate (*MeJA*) and *ABA* can cause significant reduction in yield through their direct impacts on reproductive structures **[97,98]**. As such, the proper modulation of the pathway would be necessary to preserve yield, as depicted in the hypothetical model (Fig 12).

The proposed model of a functional *DEC*-network in LPB has the necessary components of a genetic machinery that could lead to enhanced pools of active cytokinins especially in flag leaves at the time of booting and during exposure to slow but progressive drought. Yield and yield-component data collected from the drought experiments performed for the transcriptomics studies recapitulated previously reported superior performance of LPB due to *qDTY12.1* effects (Fig 1).

### Modulation of ABA response in LPB through the *qDTY12.1* mechanism

ABA signaling is central to the first line of defense against drought but not without any costs to plant development and net productivity **[97–101]**. The prioritization of cellular resources to balance the costs of survival with net productivity may require an extensive modulation of ABA responses. The overactive transcriptomic burst at booting stage in HPB, WR and IR64 are indicative of a costly and ‘*all in’* response to drought, hence greatly perturbed cellular status. In contrast, the transcriptomic response in LPB at booting stage appeared to be more modulated or *‘tamed’* (Fig 3). In other words, more is not necessarily always better as subtle changes could go a long way. Indeed, reports in other crops also showed much fewer number of differentially expressed genes in drought-tolerant genotypes compared to more sensitive genotypes **[36,37]**. The overactive transcriptomic burst in HPB, WR and IR64 based on the directionality of transcriptome fluxes may largely be associated with ABA response.

A cursory evidence for the taming of the ABA response was illustrated by the differential expression of zeaxanthin epoxidase (ZEP*; Os04g0448900*) that catalyzes the first committed step in ABA biosynthesis via the xanthophyll cycle in plastids **[99,102,103]**. Drought-mediated upregulation or downregulation of ZEP was determined as a log2 fold-change from control values for each developmental stage **(S7 Fig)**. A specific look at the booting stage showed a −0.57 log2 decrease in *ZEP* expression in LPB with drastic expression changes evident in HPB, WR, and IR64, of 4.1 log2 increase, −3.0 log2 decrease, and 1.9 log2 increase, respectively. Interestingly, inverse trends in ZEP expression across all genotypes was evident at the vegetative and grain-filling stages. The drastic differences at booting stage suggest that LPB perhaps has the mechanism that limits ABA biosynthesis and therefore modulates ABA response mechanism more efficiently. Based on the directionality of transcriptomic fluxes, it is apparent that the ‘*taming*’ effects in LPB also extend beyond the genes involved in ABA responses.

## Materials and Methods

### Minimal comparative panel

Based on extensive genotyping and yield evaluation under progressive drought **[28,104,105]**, a minimal comparative panel illustrating the differential effects of *qDTY12.1* across genetic backgrounds was established at IRRI. This panel was comprised of the Indonesian upland cultivar Way Rarem (WR; IRGC122298) as the original donor of *qDTY12.1*, the drought-sensitive mega-variety IR64 as the recurrent parent used for backcross introgression of the *qDTY12.1* from WR, and two sibling introgression lines of IR64 carrying the *qDTY12.1* from WR (IR102784:2-42-88-2-1-2, IR102784:2-90-385-1-1-3) designated as *low-yield-penalty* (LPB) and *high-yield-penalty* (HPB) lines, respectively **[22,32]**.

### Drought experiments and tissue sample collection

Parallel replicated experiments were conducted at IRRI’s Ziegler Experiment Station in Los Banos, Laguna, Philippines (14°30′ N longitude, 121°15′ E latitude) during the 2017 wet season (WS; June to November, 2017) for the irrigated and drought conditions across the minimal comparative panel. The field experiment was an alpha-lattice design with three replicates (n = 3) and three (3) individual plants per replicate that were single-seed transplanted in the field plots after establishing for 21 days in seedling beds. Control plots were maintained in standard irrigated levy based on IRRI’s standard protocols, while the drought plots were established inside a rain-out shelter facility for drought screening next to the irrigated plots **(S2 Fig) [25,26,106–109]**. Both the irrigated and drought plots were given continuous irrigation corresponding to 5 cm standing water until thirty (30) days after planting (DAP) or 51 days after sowing, when progressive drought was initiated for the treatment group by withholding water until the end of the season. A life-saving irrigation was applied to the drought plots at the point when extensive leaf rolling was observed in order to promote survival until harvest **[34]**.

Tissue sampling was conducted on three (3) plants per replicate in both the irrigated and drought conditions. Samples were comprised of pooled flag leaves with the connected leaf sheath surrounding the developing panicle. The dates of tissue sampling were synchronized as defined by the days counted backward (***t_-1_*** = vegetative) or forward (***t_1_*** = grain-filling) from the reference time-point (***t_0_*** = booting) in order to generate developmentally comparable flag leaf transcriptomes across genotypes. At ***t_-1_***, samples were collected from three (3) plants from each genotype and experimental plot, seven (7) days after the initiation of progressive drought. At ***t_0_***, samples were collected from three (3) plants from each genotype and experimental plot, twelve (12) days prior to panicle extrusion (heading). At ***t_1_***, samples were collected fifteen (15) days after anthesis when the developing grains had milky and dough-like consistencies. All samples were collected at the same time of the day (between 8:00 AM and 10:00 AM), and were immediately frozen in liquid nitrogen. Panicle length (mm), plant height (cm), tiller number per plant, reproductive tiller number per plant, and biomass per plant were recorded from all experimental plots.

### Transcriptome analysis by RNA-Seq

Total RNA was extracted from frozen flag leaves using the miRVana™ miRNA isolation kit according to manufacturer’s protocol (Invitrogen, Carlsbad, CA). RNA from three (3) individual plants in each genotype were pooled to create a composite sample representing each replicate. Two independent RNA-Seq libraries were constructed from the pooled RNA across genotypes, developmental stages, and treatments, according to standard in-house protocols **[29]**. The indexed RNA-Seq libraries were sequenced twice in the Illumina HiSeq3000 (Oklahoma Medical Research Foundation, Norman, OK) by strand-specific and paired-end sequencing at 150-bp with 20 to 40 million sequence reads per run.

Raw RNA-Seq data was processed and assembled through the established in-house data analysis pipeline **[29]**. Sequence output from the indexed RNA-Seq libraries (PRJNA378253) was preprocessed with Cutadapt (v2.10) and mapped against the Nipponbare *RefSeq* and corresponding GFF gene models (IRGSP-1.0) using the Tophat2 (v2.1.1) and Bowtie (v2.2.8.0) [40,110,111]. Gene models were further refined using Cuffmerge and differential expression was calculated with Cuffdiff on Cufflinks (v.2.2.1) with default parameters (p-value < 0.05, FDR = 5%) **[112]**. Expression of 25,786 annotated protein-coding genes were detected across the RNA-Seq data matrices of the irrigated versus drought-stressed plants at vegetative (V7 to V10), early booting (R1 to R2), and grain filling (R7) stages. Transcript abundance for each annotated locus was expressed as Fragments per Kilobase of Transcript per Million (FPKM). Biological interrogation of the transcriptome was performed in three windows, *i.e.,* global or total transcriptome (n = 25,786 loci), transcription factor genes (n = 1,340 loci), and stress-related genes (n = 2,589 loci). Transcription factors were extracted from the Nipponbare *RefSeq* (https://www.ncbi.nlm.nih.gov/genome/annotation_euk/Oryza_sativa_Japonica_Grou p/102/). Stress-related loci were extracted using the keywords listed in **S2 Table**.

### Propensity transformation of RNA-Seq data

Direct comparison of FPKM-based expression has proven to be less efficient in extracting biologically meaningful expression patterns (fluxes) because of the confounding effects created by the highly disparate nature of inter-genotypic variation and the stochastic nature of gene expression. Meaningful changes in gene expression are also dependent on the molecular interactions of target genes and their activators/repressors **[113,114]**. The Gene Flux Theory posits that within the natural competition for transcriptional machinery, genes with low transcript abundances are ultra-sensitive to the effects of other genes **[115]**. Critical loci with low FPKM in one genotype are often discarded, making the directional character of expression fluxes difficult to extract. To address these potential limitations in interpreting the biological significance of inter-genotypic differences, we performed an additional normalization of the total dataset by Propensity Transformation, which uses ‘*within genotype’* and ‘*within treatment*’ comparisons of FPKM-based expression for each locus against the summation across all time-points and against the summation of all loci across the entire dataset **[116]**. The FPKM values across the entire transcriptome matrix were Propensity-transformed and normalized by: 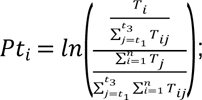 Where: *Pt_i_* = Propensity transformation of FPKM of transcript *I; T_i_* = FPKM of transcript *I; n* = Total number of transcripts (25,786); *j* = Variable that iterates over datasets of *t_1_* =vegetative, *t_2_* =booting, *t_3_* = grain-filling; and *i* = Variable that iterates over the total number of transcript-encoding loci **(S3 Fig)**. Propensity-transformed datasets (global, transcription factor, and stress-related windows) were filtered ata threshold of *-0.3<PROPENSITY>0.3* in order to extract the gene loci with the largest fluxes, hence most biologically informative differences. The total of 8,215 loci (out of 25,786) from the global dataset were subjected to k-means clustering to further refine the large cohort into fifteen (15) sub-clusters for Propensity ≥ 0.3, and ten (10) sub-clusters for Propensity ≤ −0.3. One sub-cluster was chosen from the extremes of each group and the loci were combined into 384 in the global filtered dataset. The transcription factor and stress-related groups did not require k-means clustering with 410 (out of 1,340) and 833 (out of 2,589) loci, respectively.

### Analysis of transcriptome fluxes and directionality

The standard approach for revealing biologically meaningful trends in RNA-Seq datasets is to identify differentially expressed genes (DEG) that correlate with the phenotype. While this approach can give useful insights into cellular processes, the underlying concept tend to be simplistic because responses at the cellular and whole organismal levels more often than not involve large number of genes **[117]**. In order to capture a more biologically relevant view of the drought response transcriptomes across genotypes, the Propensity-normalized expressions were interrogated to uncover similarities and differences in fluxes on a locus-by-locus plane. Propensity-transformed expression values facilitated direct comparison to generate profiles of expression fluxes between genotypes by hierarchical clustering of the filtered and un-filtered propensity scores in the global, transcription factor, and stress-related windows.

Analysis of the directional character of expression fluxes indicates the degree by which transcriptional responses are modulated. An unmitigated or ‘*untamed*’ transcriptional response would be characterized by an overabundance of positive transcriptional activities while a modulated or ‘*tamed*’ response would be characterized by highly regulated or controlled repression. The directional character of expression fluxes in the unfiltered dataset was assessed by comparing the fraction of loci with positive propensity scores (PPF – positive propensity fraction) to the fraction of loci with negative propensity scores (NPF – negative propensity fraction). Directionality was scored as positive skew (upward pointing arrow), negative skew (downward pointing arrow), or neutral (line segment) (Fig 3). A positive skew was given when PPF was greater than NPF, while a negative skew was given when NPF was greater than PPF, and neutral when PPF was approximately equal to NPF. Genes with Propensity scores = 0 (5% to 9% of total) were excluded.

### Hierarchical clustering and statistical analysis

Hierarchical clustering and other statistical analyses were performed using JMP® (v14.0.0. SAS Institute Inc., Cary, NC). Mean comparisons of agronomic measurements in rice and Arabidopsis experiments were performed with Tukey HSD following a significant analysis of variance at p = 0.05.

### RiceFREND and KnetMiner analyses

The RiceFREND online analysis portal was used for initial capture of other genes that are co-expressed with *OsDEC* (https://ricefrend.dna.affrc.go.jp/) **[41]**. The multiple gene guide tool was used to determine co-expression of eighteen (18) expressed genes of *qDTY12.1* to generate a co-expression map by default setting.

The KnetMiner tool was used determine the enrichment of biochemical, physiological, and agronomic traits that are associated with the various components of the *DEC*-network (http://knetminer.rothamsted.ac.uk) **[55]**. The Rap-DB loci for the 36 genes in the *DEC*-network was used as the query to search for domains (relevant biological processes) in Knetminer using default parameters **[40]**. This tool integrates knowledge in public domain related to the query (*e.g.,* gene function, GWAS, Protein, Phenotype, Pathways, etc.) to generate a knowledge map of biological functions.

### Analysis of *DEC* knock-out mutants in Arabidopsis

Orthologs of *OsDEC* in *Arabidopsis thaliana* were determined as *At3G03460* (3G^m^) and *At5G17510* (5Gm) **[33]**, and confirmed by phylogenetic analysis with EnsemblPlants (https://plants.ensembl.org/Arabidopsis_thaliana/Info/Index). Homozygous mutants **[118]** were determined using the Salk Institute TDNA Express Gene Mapping Tools (http://signal.salk.edu/cgi-bin/tdnaexpress) and seeds were obtained from the Arabidopsis Biological Research Center (https://abrc.osu.edu/). Seeds were vernalized in 0.1% (w/v) agarose at 4°C for 7 days, sown onto moistened peat pellets (Jiffy-7® – Peat Pellets) and grown for fourteen (14) days in growth chambers (Percival Scientific) at constant 22°C with 16 hours of light (100 µmol m^-2^ s^-1^) and 60-70% relative humidity. DNA and RNA extraction and PCR and qRT-PCR analyses of the mutants were performed according to standard protocols using the primer sets listed in **S6 Table**.

The agronomic and yield performances of *AtDEC* wild-type and mutants were investigated in a growth chamber drought experiments that mirrored the developmental timing of stress in the rice experiments **[119]**. A pilot study established the effective drought conditions at 30% field capacity, eight (8) days prior to bolting, 27°C day/22°C night, and 40% relative humidity. Control experiments were performed at 70% field capacity, constant 22°C, and 65-80% relative humidity. Vernalized seeds of Col-0, 3G^m^, and 5G^m^ were germinated in peat pellets (Jiffy-7^®^ Peat Pellets, 42mm x 65mm) and grown in two separate growth chambers at constant 22°C with 16 hours of light (100 µmol m^-2^ s^-1^) and 65-80% relative humidity. Drought experiment was performed by growing the plants for 20 days (Col-0; 3G^m^) and 16 days (5G^m^), before withholding irrigation. Progressive drought was imposed for 14 days by maintaining 30% field capacity, while the control plants were maintained at 70% field capacity. The peat pellets at 30% field capacity received a daily water input to maintain a weight of 33 g (peat pellet + plant + plus water input). Control and post-drought plants were maintained at 65-70 g. Days to bolting, days to first bloom, days to seed set (first silique), total dry biomass per plant, and total seed yield per plant were determined under both irrigated and drought conditions.

## Acknowledgments

This project was supported by NSF-IOS Plant Genome Research Program Grant-1602494 and Bayer CropScience Endowed Professorship. Genomic computations were performed using the supercomputing facilities at the ROIS National Institute of Genetics, Mishima, Japan, and Texas Tech University High-Performance Computing Cluster. Next-Gen sequencing was performed at the Oklahoma Medical Research Foundation, Norman, OK.

## Supporting information

**S1 Fig.** Diagrammatic representation of the recombination dynamics that likely occurred in the breeding scheme that generated the two *qDTY12.1* sibling introgression lines (LPB, HPB). The divergent phenotypes exhibited by LPB and HPB are enigmatic considering that both carry the *qDTY12.1* introgression and both were derived from the same parental lineages (Way Rarem, Vandana, and IR64) via backcross coupled with marker assisted selection. LPB and HPB each underwent separate and distinct shuffling of alleles in the genomic background, thereby creating either a synergistic (LPB) or antagonistic (HPB) *DEC*-network.

**S2 Fig.** Schematic of the drought experiment conducted at IRRI in the wet season of 2017 under a rain-shelter facility. Rice plants were exposed to progressive drought by withholding irrigation at the vegetative stage through maturity with only a single life-saving irrigation applied after the initiation of stress. Flag leaves were collected at the vegetative, booting, and grain-filling stages across the comparative panel to generate the RNASeq transcriptomic data data matrix. Definition of developmental stages were according to current standards **[68]**.

**S3 Fig.** Distribution plots of Propensity scores calculated from the FPKM-based expression values across the flag leaf RNASeq transcriptomic datasets. Propensity score distributions of transcript abundance (FPKM) for 25,786 loci at (**A**) vegetative, (**B**) booting, (**C**), and grain-fill stages under control/irrigated conditions, and (**D**) vegetative, (**E**) booting, and (**F**) grain-filling stages under stress/drought conditions.

**S4 Fig.** Phylogenetic analysis of *OsDEC* (Os12g0465700). Homology of the rice *DECUSSATE* gene to *Arabidopsis thaliana* At3G03460 and At5G17510 orthologs was referenced in the published report **[33]**, and re-validated by using the phylogenetic tree function at EnsemblPlants (https://plants.ensembl.org/index.html).

**S5 Fig.** Comparison of reproductive milestones of the *Arabidopsis thaliana AtDEC* T-DNA insertion mutants At3g03460 (3Gm), At5G17510 (5Gm) and Columbia wild-type (Col-0) under control and drought conditions. (**A**) Days to bolting; (**B**) Days to first bloom; (**C**) Days to seed set. Box plots are means of individual plants (n = 16). Means were separated with Tukey HSD after significant ANOVA. **significant difference at at p<0.01; ***significant difference at p<0.001; n.s. – no significant difference at p<0.05.

**S6 Fig.** Expression of *OsDST* (Os03g0786400; Drought and Salt Tolerance) during (**A**) Vegetative, (**B**) Booting, (**C**) Grain-filling stages. *OsDST* directly regulates cytokinin degradation via *OsCKX2*, which is a single copy gene in the rice genome. During the critical stage of booting, *OsDST* was downregulated LPB and IR64 with - 2.24 and −5.86 log2 fold-change, respectively. *OsDST* was upregulated in HPB and WR at 0.74 and 0.62 log2 fold-change, respectively.

**S7 Fig.** Expression of *OsZEP* (Os04g0448900; zeathanthin expoxidase) during (**A**) Vegetative, (**B**) Booting, and (**C**) Grain-filling stages. Involved in the first committed step of ABA biosynthesis, *OsZEP* activity directly impacts the ABA response, especially under abiotic stress. At booting stage, LPB and WR exhibited decreases in *OsZEP* expression with −0.57 and −3.0 log2 fold-change, respectively. In contrast, HPB and WR exhibited increases with 4.1 and 1.9 log2 fold-change, respectively.

**S1 Table.** List of qDTYs that are known to contribute to yield retention under reproductive-stage drought.

**S2 Table.** Key terms used to extract stress-related genes from the global transcriptomic window of 25,786 loci.

**S3 Table.** List of annotated protein-coding gene loci (n = 50) within the *qDTY12.1* boundaries.

**S4 Table.** List of the genes that comprised the full DEC-network (n = 36) that is operative during the onset of booting under drought.

**S5 Table.** Biological functions and traits that were associated to *DEC*-network by the KnetMiner knowledge integration platform.

**S6 Table.** DNA primers and methods used for genomic-PCR (genotyping) and qRT-PCR analyses of Arabidopsis *AtDEC* T-DNA insertion mutants.

## References

1. Khush GS. Strategies for increasing the yield potential of cereals: case of rice as an example. Gupta P, editor. Plant Breed. 2013;132: n/a-n/a. doi:10.1111/pbr.1991

2. Peng S, Khush GS, Virk P, Tang Q, Zou Y. Progress in ideotype breeding to increase rice yield potential. F Crop Res. 2008;108: 32–38. doi:10.1016/j.fcr.2008.04.001

3. Khush GS. What it will take to Feed 5.0 Billion Rice consumers in 2030. Plant Mol Biol. 2005;59: 1–6. doi:10.1007/s11103-005-2159-5

4. Khush GS. Green revolution: the way forward. Nat Rev Genet. 2001;2: 815–822. doi:10.1038/35093585

5. Khush GS. Modern varieties — Their real contribution to food supply and equity. GeoJournal. 1995;35: 275–284. doi:10.1007/BF00989135

6. Khush GS. Breaking the yield frontier of rice. GeoJournal. 1995;35: 329–332. doi:10.1007/BF00989140

7. de los Reyes BG. Genomic and epigenomic bases of transgressive segregation – New breeding paradigm for novel plant phenotypes. Plant Sci. 2019;288: 110213. doi:10.1016/j.plantsci.2019.110213

8. Pabuayon ICM, Kitazumi A, Gregorio GB, Singh RK, de los Reyes BG. Contributions of Adaptive Plant Architecture to Transgressive Salinity Tolerance in Recombinant Inbred Lines of Rice: Molecular Mechanisms Based on Transcriptional Networks. Frontiers in Genetics. 2020. p. 1318. Available: https://www.frontiersin.org/article/10.3389/fgene.2020.594569

9. Sandhu N, Kumar A. Bridging the Rice Yield Gaps under Drought: QTLs, Genes, and their Use in Breeding Programs. Agronomy. 2017;7: 27. doi:10.3390/agronomy7020027

10. Dar MH, Waza SA, Shukla S, Zaidi NW, Nayak S, Hossain M, et al. Drought Tolerant Rice for Ensuring Food Security in Eastern India. Sustainability. 2020;12: 2214. doi:10.3390/su12062214

11. Lei Y, Liu C, Zhang L, Luo S. How smallholder farmers adapt to agricultural drought in a changing climate: A case study in southern China. Land use policy. 2016;55: 300– 308. doi:10.1016/j.landusepol.2016.04.012

12. Serraj R, McNally KL, Slamet-Loedin I, Kohli A, Haefele SM, Atlin G, et al. Drought Resistance Improvement in Rice: An Integrated Genetic and Resource Management Strategy. Plant Prod Sci. 2011;14: 1–14. doi:10.1626/pps.14.1

13. Lottering S, Mafongoya P, Lottering R. Drought and its impacts on small-scale farmers in sub-Saharan Africa: a review. South African Geogr J. 2020; 1–23. doi:10.1080/03736245.2020.1795914

14. Vikram P, Swamy BPMBM, Dixit S, Ahmed HU, Teresa Sta Cruz M, Singh AK, et al. QDTY 1.1, a major QTL for rice grain yield under reproductive-stage drought stress with a consistent effect in multiple elite genetic backgrounds. BMC Genet. 2011;12. doi:10.1186/1471-2156-12-89

15. Singh R, Singh Y, Xalaxo S, Verulkar S, Yadav N, Singh S, et al. From QTL to variety-harnessing the benefits of QTLs for drought, flood and salt tolerance in mega rice varieties of India through a multi-institutional network. Plant Sci. 2016;242: 278–287. doi:10.1016/j.plantsci.2015.08.008

16. Swamy BPM, Kumar A, Cruz PCS, Shamsudin NAA, Boromeo TH, Palanog AD, et al. Grain yield QTLs with consistent-effect under reproductive-stage drought stress in rice. F Crop Res. 2014;161: 46–54. doi:10.1016/j.fcr.2014.01.004

17. Bernier J, Kumar A, Ramaiah V, Spaner D, Atlin G. A large-effect QTL for grain yield under reproductive-stage drought stress in upland rice. Crop Sci. 2007;47: 507–518. doi:10.2135/cropsci2006.07.0495

18. Bernier J, Kumar A, Venuprasad R, Spaner D, Verulkar S, Mandal NP, et al. Characterization of the effect of a QTL for drought resistance in rice, qtl12.1, over a range of environments in the Philippines and eastern India. Euphytica. 2009;166: 207– 217. doi:10.1007/s10681-008-9826-y

19. Mishra KK, Vikram P, Yadaw RB, Swamy BM, Dixit S, Cruz MTS, et al. qDTY12.1: a locus with a consistent effect on grain yield under drought in rice. BMC Genet. 2013;14: 12. doi:10.1186/1471-2156-14-12

20. Dixit S, Swamy BPM, Vikram P, Ahmed HU, Sta Cruz MT, Amante M, et al. Fine mapping of QTLs for rice grain yield under drought reveals sub-QTLs conferring a response to variable drought severities. Theor Appl Genet. 2012;125: 155–169. doi:10.1007/s00122-012-1823-9

21. Swamy BPM, Vikram P, Dixit S, Ahmed HU, Kumar A. Meta-analysis of grain yield QTL identified during agricultural drought in grasses showed consensus. BMC Genomics. 2011;12: 319. doi:10.1186/1471-2164-12-319

22. Yadav S, Sandhu N, Majumder RR, Dixit S, Kumar S, Singh SP, et al. Epistatic interactions of major effect drought QTLs with genetic background loci determine grain yield of rice under drought stress. Sci Rep. 2019;9: 2616. doi:10.1038/s41598-019-39084-7

23. Dixit S, Kumar Biswal A, Min A, Henry A, Oane RH, Raorane ML, et al. Action of multiple intra-QTL genes concerted around a co-localized transcription factor underpins a large effect QTL. Sci Rep. 2015;5. doi:10.1038/srep15183

24. Henry A, Stuart-Williams H, Dixit S, Kumar A, Farquhar G. Stomatal conductance responses to evaporative demand conferred by rice drought-yield quantitative trait locus qDTY12.1. Funct Plant Biol. 2019;46: 660. doi:10.1071/FP18126

25. Henry A, Swamy BPM, Dixit S, Torres RD, Batoto TC, Manalili M, et al. Physiological mechanisms contributing to the QTL-combination effects on improved performance of IR64 rice NILs under drought. J Exp Bot. 2015;66: 1787–1799. doi:10.1093/jxb/eru506

26. Henry A, Dixit S, Mandal NP, Anantha MS, Torres R, Kumar A. Grain yield and physiological traits of rice lines with the drought yield QTL qDTY12.1 showed different responses to drought and soil characteristics in upland environments. Funct Plant Biol. 2014;41: 1066. doi:10.1071/FP13324

27. Raorane ML, Pabuayon IM, Miro B, Kalladan R, Reza-Hajirezai M, Oane RH, et al. Variation in primary metabolites in parental and near-isogenic lines of the QTL qDTY12.1: altered roots and flag leaves but similar spikelets of rice under drought. Mol Breed. 2015;35: 1–25. doi:10.1007/s11032-015-0322-5

28. Kumar A, Dixit S, Ram T, Yadaw RB, Mishra KK, Mandal NP. Breeding high-yielding drought-tolerant rice: genetic variations and conventional and molecular approaches. J Exp Bot. 2014;65: 6265–6278. doi:10.1093/jxb/eru363

29. Kitazumi A, Pabuayon ICM, Ohyanagi H, Fujita M, Osti B, Shenton MR, et al. Potential of Oryza officinalis to augment the cold tolerance genetic mechanisms of Oryza sativa by network complementation. Sci Rep. 2018;8: 16346. doi:10.1038/s41598-018-34608-z

30. Pabuayon ICM, Kitazumi A, Cushman KR, Singh RK, Gregorio GB, Dhatt BK, et al. Novel and transgressive salinity tolerance in recombinant inbred lines of rice created by physiological coupling-uncoupling and network rewiring effects. Front Plant Sci. 2021. doi:10.3389/fpls.2021.615277

31. Boyle EA, Li YI, Pritchard JK. An Expanded View of Complex Traits: From Polygenic to Omnigenic. Cell. 2017;169: 1177–1186. doi:10.1016/j.cell.2017.05.038

32. Kumar A, Sandhu N, Venkateshwarlu C, Priyadarshi R, Yadav S, Majumder RR, et al. Development of introgression lines in high yielding, semi-dwarf genetic backgrounds to enable improvement of modern rice varieties for tolerance to multiple abiotic stresses free from undesirable linkage drag. Sci Rep. 2020;10: 13073. doi:10.1038/s41598-020-70132-9

33. Itoh J, Hibara K, Kojima M, Sakakibara H, Nagato Y. Rice DECUSSATE controls phyllotaxy by affecting the cytokinin signaling pathway. Plant J. 2012;72: 869–881. doi:10.1111/j.1365-313x.2012.05123.x

34. Torres R, Henry A, Kumar A. Methodologies for managed drought stress experiments in the field. In: Shashidhar HE, Henry A, Hardy B, editors. Methodologies for root drought studies in rice. Los Banos, Phillipines: International Rice Research Institute; 2012. pp. 43–50.

35. Bechtold U, Field B. Molecular mechanisms controlling plant growth during abiotic stress. J Exp Bot. 2018;69: 2753–2758. doi:10.1093/jxb/ery157

36. Fracasso A, Trindade LM, Amaducci S. Drought stress tolerance strategies revealed by RNA-Seq in two sorghum genotypes with contrasting WUE. BMC Plant Biol. 2016;16: 115. doi:10.1186/s12870-016-0800-x

37. Bhogireddy S, Xavier A, Garg V, Layland N, Arias R, Payton P, et al. Genome-wide transcriptome and physiological analyses provide new insights into peanut drought response mechanisms. Sci Rep. 2020;10: 4071. doi:10.1038/s41598-020-60187-z

38. Boonjung H, Fukai S. Effects of soil water deficit at different growth stages on rice growth and yield under upland conditions. 2. Phenology, biomass production and yield. F Crop Res. 1996;48: 47–55. doi:https://doi.org/10.1016/0378-4290(96)00039-1

39. Zhang J, Zhang S, Cheng M, Jiang H, Zhang X, Peng C, et al. Effect of Drought on Agronomic Traits of Rice and Wheat: A Meta-Analysis. Int J Environ Res Public Health. 2018;15: 839. doi:10.3390/ijerph15050839

40. Sakai H, Lee SS, Tanaka T, Numa H, Kim J, Kawahara Y, et al. Rice Annotation Project Database (RAP-DB): An Integrative and Interactive Database for Rice Genomics. Plant Cell Physiol. 2013;54: e6–e6. doi:10.1093/pcp/pcs183

41. Sato Y, Namiki N, Takehisa H, Kamatsuki K, Minami H, Ikawa H, et al. RiceFREND: a platform for retrieving coexpressed gene networks in rice. Nucleic Acids Res. 2012/11/23. 2013;41: D1214–D1221. doi:10.1093/nar/gks1122

42. Andersen SU, Algreen-Petersen RG, Hoedl M, Jurkiewicz A, Cvitanich C, Braunschweig U, et al. The conserved cysteine-rich domain of a tesmin/TSO1-like protein binds zinc in vitro and TSO1 is required for both male and female fertility in Arabidopsis thaliana. J Exp Bot. 2007;58: 3657–3670. doi:10.1093/jxb/erm215

43. Wang Y, Zhang W-Z, Song L-F, Zou J-J, Su Z, Wu W-H. Transcriptome Analyses Show Changes in Gene Expression to Accompany Pollen Germination and Tube Growth in Arabidopsis. Plant Physiol. 2008;148: 1201–1211. doi:10.1104/pp.108.126375

44. Hauser BA, He JQ, Park SO, Gasser CS. TSO1 is a novel protein that modulates cytokinesis and cell expansion in Arabidopsis. Development. 2000;127: 2219–2226. doi:10.1590/0104-07072017005650015

45. Sijacic P, Wang W, Liu Z. Recessive Antimorphic Alleles Overcome Functionally Redundant Loci to Reveal TSO1 Function in Arabidopsis Flowers and Meristems. Qu L-J, editor. PLoS Genet. 2011;7: e1002352. doi:10.1371/journal.pgen.1002352

46. Klepikova A V., Kasianov AS, Gerasimov ES, Logacheva MD, Penin AA. A high resolution map of the Arabidopsis thaliana developmental transcriptome based on RNA-seq profiling. Plant J. 2016;88: 1058–1070. doi:10.1111/tpj.13312

47. Reguera M, Peleg Z, Abdel-Tawab YM, Tumimbang EB, Delatorre CA, Blumwald E. Stress-Induced Cytokinin Synthesis Increases Drought Tolerance through the Coordinated Regulation of Carbon and Nitrogen Assimilation in Rice. PLANT Physiol. 2013;163: 1609–1622. doi:10.1104/pp.113.227702

48. D’Aloia M, Bonhomme D, Bouché F, Tamseddak K, Ormenese S, Torti S, et al. Cytokinin promotes flowering of Arabidopsis via transcriptional activation of the FT paralogue TSF. Plant J. 2011;65: 972–979. doi:10.1111/j.1365-313X.2011.04482.x

49. Bartrina I, Otto E, Strnad M, Werner T, Schmülling T. Cytokinin Regulates the Activity of Reproductive Meristems, Flower Organ Size, Ovule Formation, and Thus Seed Yield in Arabidopsis thaliana. Plant Cell. 2011;23: 69–80. doi:10.1105/tpc.110.079079

50. Murai N. Review: Plant Growth Hormone Cytokinins Control the Crop Seed Yield. Am J Plant Sci. 2014;05: 2178–2187. doi:10.4236/ajps.2014.514231

51. Zahir ZA, Asghar HN, Arshad M. Cytokinin and its precursors for improving growth and yield of rice. Soil Biol Biochem. 2001;33: 405–408. doi:10.1016/S0038-0717(00)00145-0

52. Jameson PE, Song J. Cytokinin: a key driver of seed yield. J Exp Bot. 2016;67: 593– 606. doi:10.1093/jxb/erv461

53. Wang M, Lu X, Xu G, Yin X, Cui Y, Huang L, et al. OsSGL, a novel pleiotropic stress-related gene enhances grain length and yield in rice. Sci Rep. 2016;6: 38157. doi:10.1038/srep38157

54. Inukai Y, Nagato Y, Nonomura K-I, Kitano H, Itoh J-I, Yamaki S, et al. Rice Plant Development: from Zygote to Spikelet. Plant Cell Physiol. 2005;46: 23–47. doi:10.1093/pcp/pci501

55. Hassani-Pak K. KnetMiner - An integrated data platform for gene mining and biological knowledge discovery. Universität Bielefeld. 2017.

56. Wu J, Lawit SJ, Weers B, Sun J, Mongar N, Van Hemert J, et al. Overexpression of zmm28 increases maize grain yield in the field. Proc Natl Acad Sci. 2019/11/04. 2019;116: 23850–23858. doi:10.1073/pnas.1902593116

57. Becker A, Theißen G. The major clades of MADS-box genes and their role in the development and evolution of flowering plants. Mol Phylogenet Evol. 2003;29: 464– 489. doi:https://doi.org/10.1016/S1055-7903(03)00207-0

58. Ng M, Yanofsky MF. FUNCTION AND EVOLUTION OF THE PLANT MADS-BOX GENE FAMILY. Nat Rev Genet. 2001;2: 186–195. doi:http://dx.doi.org/10.1038/35056041

59. Kater MM, Dreni L, Colombo L. Functional conservation of MADS-box factors controlling floral organ identity in rice and Arabidopsis. J Exp Bot. 2006;57: 3433– 3444. doi:10.1093/jxb/erl097

60. Ghimire KH, Quiatchon LA, Vikram P, Swamy BPM, Dixit S, Ahmed H, et al. Identification and mapping of a QTL (qDTY1.1) with a consistent effect on grain yield under drought. F Crop Res. 2012;131: 88–96. doi:https://doi.org/10.1016/j.fcr.2012.02.028

61. Sandhu N, Singh A, Dixit S, Sta Cruz MT, Maturan PC, Jain RK, et al. Identification and mapping of stable QTL with main and epistasis effect on rice grain yield under upland drought stress. BMC Genet. 2014;15: 1–15. doi:10.1186/1471-2156-15-63

62. Vikram P, Swamy BPMM, Dixit S, Trinidad J, Cruz MTS, Maturan PC, et al. Linkages and Interactions Analysis of Major Effect Drought Grain Yield QTLs in Rice. PLoS One. 2016;11: e0151532. doi:10.1371/journal.pone.0151532

63. Han J-H, Shin N-H, Moon J-H, Yi C, Yoo S-C, Chin JH. Genetic and Phenotypic Characterization of Rice Backcrossed Inbred Sister Lines of Saltol in Temperate Saline Reclaimed Area. Plant Breed Biotechnol. 2020/03/01. 2020;8: 58–68. doi:10.9787/PBB.2020.8.1.58

64. Rahman MA, Haque ME, Sikdar B, Islam MA, Matin M. Correlation Analysis of Flag Leaf with Yield in Several Rice Cultivars. J Life Earth Sci. 2014;8. doi:10.3329/jles.v8i0.20139

65. Abdalla Basyouni Abou-Khalifa A, Misra AN, El-Azeem M Salem AK. Effect of leaf cutting on physiological traits and yield of two rice cultivars. African J Plant Sci. 2008;2: 147–150. Available: http://www.academicjournals.org/AJPS

66. Cui K, Peng S, Xing Y, Yu S, Xu C, Zhang Q. Molecular dissection of the genetic relationships of source, sink and transport tissue with yield traits in rice. Theor Appl Genet. 2003;106: 649–658. doi:10.1007/s00122-002-1113-z

67. Yoshida S. Fundamentals of Rice Crop Sciene. Los Banos, Philippines: International Rice Research Institute; 1981.

68. Counce PA, Keisling TC, Mitchell AJ. A Uniform, Objective, and Adaptive System for Expressing Rice Development. Crop Sci. 2000;40: 436–443. doi:https://doi.org/10.2135/cropsci2000.402436x

69. Torres RO, Henry A. Yield stability of selected rice breeding lines and donors across conditions of mild to moderately severe drought stress. F Crop Res. 2018;220. doi:10.1016/j.fcr.2016.09.011

70. To JPC, Haberer G, Ferreira FJ, Deruère J, Mason MG, Schaller GE, et al. Type-A Arabidopsis Response Regulators Are Partially Redundant Negative Regulators of Cytokinin Signaling. Plant Cell. 2004/02/18. 2004;16: 658–671. doi:10.1105/tpc.018978

71. Hill K, Mathews DE, Kim HJ, Street IH, Wildes SL, Chiang Y-H, et al. Functional Characterization of Type-B Response Regulators in the Arabidopsis Cytokinin Response. Plant Physiol. 2013;162: 212 LP – 224. doi:10.1104/pp.112.208736

72. Xie M, Chen H, Huang L, O’Neil RC, Shokhirev MN, Ecker JR. A B-ARR-mediated cytokinin transcriptional network directs hormone cross-regulation and shoot development. Nat Commun. 2018;9: 1–13. doi:10.1038/s41467-018-03921-6

73. Ashikari M. Cytokinin Oxidase Regulates Rice Grain Production. Science (80-). 2005;309: 741–745. doi:10.1126/science.1113373

74. Peleg Z, Reguera M, Tumimbang E, Walia H, Blumwald E. Cytokinin-mediated source/sink modifications improve drought tolerance and increase grain yield in rice under water-stress. Plant Biotechnol J. 2011;9: 747–758. doi:10.1111/j.1467-7652.2010.00584.x

75. Qin H, Zhang Y, Sun L, Gu Q, Kuppu S, Zhang H, et al. Regulated Expression of an Isopentenyltransferase Gene (IPT) in Peanut Significantly Improves Drought Tolerance and Increases Yield Under Field Conditions. Plant Cell Physiol. 2011;52: 1904–1914. doi:10.1093/pcp/pcr125

76. Zhu X, Sun L, Kuppu S, Hu R, Mishra N, Smith J, et al. The yield difference between wild-type cotton and transgenic cotton that expresses IPT depends on when water-deficit stress is applied. Sci Rep. 2018;8: 2538. doi:10.1038/s41598-018-20944-7

77. Kuppu S, Mishra N, Hu R, Sun L, Zhu X, Shen G, et al. Water-deficit inducible expression of a cytokinin biosynthetic gene IPT improves drought tolerance in cotton. PLoS One. 2013;8: e64190–e64190. doi:10.1371/journal.pone.0064190

78. Henry A, Stuart-Williams B H, Dixit S, Kumar A, Farquhar G. Stomatal conductance responses to evaporative demand conferred by rice drought-yield quantitative trait locus qDTY 12.1. doi:10.1071/FP18126

79. Raorane ML, Pabuayon IM, Varadarajan AR, Mutte SK, Kumar A, Treumann A, et al. Proteomic insights into the role of the large-effect QTL qDTY 12.1 for rice yield under drought. Mol Breed. 2015;35: 139. doi:10.1007/s11032-015-0321-6

80. Lee YS, An G. Regulation of flowering time in rice. J Plant Biol. 2015;58: 353–360. doi:10.1007/s12374-015-0425-x

81. Fornara F, Pařenicová L, Falasca G, Pelucchi N, Masiero S, Ciannamea S, et al. Functional Characterization of OsMADS18, a Member of the AP1/SQUA Subfamily of MADS Box Genes. Plant Physiol. 2004;135: 2207–2219. Available: http://www.jstor.org/stable/4356576

82. Kumar A, Verulkar S, Dixit S, Chauhan B, Bernier J, Venuprasad R, et al. Yield and yield-attributing traits of rice (Oryza sativa L.) under lowland drought and suitability of early vigor as a selection criterion. F Crop Res. 2009;114: 99–107. doi:10.1016/j.fcr.2009.07.010

83. Pantuwan G, Fukai S, Cooper M, Rajatasereekul S, O’Toole JC. Yield response of rice (Oryza sativa L.) genotypes to different types of drought under rainfed lowlands. Part Grain yield and yield components. F Crop Res. 2002;73: 153–168.

84. Saluja D, Vassallo MF, Tanese N. Distinct Subdomains of Human TAFII130 Are Required for Interactions with Glutamine-Rich Transcriptional Activators. Mol Cell Biol. 1998;18: 5734–5743. doi:10.1128/MCB.18.10.5734

85. Freiman RN, Tjian R. A Glutamine-Rich Trail Leads to Transcription Factors. Science (80-). 2002;296: 2149 LP – 2150. doi:10.1126/science.1073845

86. Ding Y-H, Liu N-Y, Tang Z-S, Liu J, Yang W-C. Arabidopsis GLUTAMINE-RICH PROTEIN23 Is Essential for Early Embryogenesis and Encodes a Novel Nuclear PPR Motif Protein That Interacts with RNA Polymerase II Subunit III. Plant Cell. 2006/02/17. 2006;18: 815–830. doi:10.1105/tpc.105.039495

87. Rahman S, Sowa ME, Ottinger M, Smith JA, Shi Y, Harper JW, et al. The Brd4 Extraterminal Domain Confers Transcription Activation Independent of pTEFb by Recruiting Multiple Proteins, Including NSD3. Mol Cell Biol. 2011;31: 2641–2652. doi:10.1128/MCB.01341-10

88. Wu S-Y, Chiang C-M. The Double Bromodomain-containing Chromatin Adaptor Brd4 and Transcriptional Regulation. J Biol Chem. 2007;282: 13141–13145. doi:10.1074/jbc.R700001200

89. Huang X-Y, Chao D-Y, Gao J-P, Zhu M-Z, Shi M, Lin H-X. A previously unknown zinc finger protein, DST, regulates drought and salt tolerance in rice via stomatal aperture control. Genes Dev. 2009;23: 1805–1817. doi:10.1101/gad.1812409

90. Li S, Zhao B, Yuan D, Duan M, Qian Q, Tang L, et al. Rice zinc finger protein DST enhances grain production through controlling Gn1a/OsCKX2 expression. Proc Natl Acad Sci. 2013;110: 3167–3172. doi:10.1073/pnas.1300359110

91. Zürcher E, Müller B. Cytokinin Synthesis, Signaling, and Function—Advances and New Insights. In: Jeon KWBT-IR of C and MB, editor. International Review of Cell and Molecular Biology. Academic Press; 2016. pp. 1–38. doi:https://doi.org/10.1016/bs.ircmb.2016.01.001

92. Hwang I, Sheen J, Müller B. Cytokinin Signaling Networks. Annu Rev Plant Biol. 2012;63: 353–380. doi:10.1146/annurev-arplant-042811-105503

93. El-Showk S, Ruonala R, Helariutta Y. Crossing paths: Cytokinin signalling and crosstalk. Dev. 2013;140: 1373–1383. doi:10.1242/dev.086371

94. Kim SL, Lee S, Kim HJ, Nam HG, An G. OsMADS51 is a short-day flowering promoter that functions upstream of Ehd1, OsMADS14, and Hd3a. Plant Physiol. 2007/10/19. 2007;145: 1484–1494. doi:10.1104/pp.107.103291

95. Weng X, Wang L, Wang J, Hu Y, Du H, Xu C, et al. Grain Number, Plant Height, and Heading Date7 is a central regulator of growth, development, and stress response. Plant Physiol. 2014;164: 735–747. doi:10.1104/pp.113.231308

96. Yoshida H, Nagato Y. Flower development in rice. J Exp Bot. 2011;62: 4719–4730. doi:10.1093/jxb/err272

97. Davies WJ, Wilkinson S, Veselov DS, Kudoyarova GR, Arkhipova TN. Plant hormone interactions: innovative targets for crop breeding and management. J Exp Bot. 2012;63: 3499–3509. doi:10.1093/jxb/ers148

98. Kim EH, Kim YS, Park SH, Koo YJ, Choi Y Do, Chung YY, et al. Methyl jasmonate reduces grain yield by mediating stress signals to alter spikelet development in rice. Plant Physiol. 2009;149: 1751–1760. doi:10.1104/pp.108.134684

99. Tuteja N. Abscisic Acid and Abiotic Stress Signaling. Plant Signal Behav. 2007;2: 135–138. doi:10.4161/psb.2.3.4156

100. Finkelstein RR, Rock CD. Abscisic Acid Biosynthesis and Response. Arab B. 2002;2002. doi:10.1199/tab.0058

101. Zhang J, Jia W, Yang J, Ismail AM. Role of ABA in integrating plant responses to drought and salt stresses. F Crop Res. 2006;97: 111–119. doi:10.1016/j.fcr.2005.08.018

102. Taylor IB, Burbidge A, Thompson AJ. Control of abscisic acid synthesis. J Exp Bot. 2000;51: 1563–1574. doi:10.1093/jexbot/51.350.1563

103. Verma V, Ravindran P, Kumar PP. Plant hormone-mediated regulation of stress responses. BMC Plant Biol. 2016;16: 86. doi:10.1186/s12870-016-0771-y

104. Kumar A, Bernier J, Verulkar S, Lafitte HR, Atlin GN. Breeding for drought tolerance: Direct selection for yield, response to selection and use of drought-tolerant donors in upland and lowland-adapted populations. F Crop Res. 2008;107: 221–231. doi:10.1016/j.fcr.2008.02.007

105. Dixit S, Singh A, Kumar A. Rice breeding for high grain yield under drought: A strategic solution to a complex problem. Int J Agron. 2014;2014. doi:10.1155/2014/863683

106. Villa JE, Henry A, Xie F, Serraj R. Hybrid rice performance in environments of increasing drought severity. F Crop Res. 2012;125: 14–24. doi:10.1016/j.fcr.2011.08.009

107. Torres RO, Henry A. Yield stability of selected rice breeding lines and donors across conditions of mild to moderately severe drought stress. F Crop Res. 2018;220. doi:10.1016/j.fcr.2016.09.011

108. Torres RO, McNally KL, Cruz CV, Serraj R, Henry A. Screening of rice Genebank germplasm for yield and selection of new drought tolerance donors. F Crop Res. 2013;147: 12–22. doi:https://doi.org/10.1016/j.fcr.2013.03.016

109. Henry A, Gowda VRP, Torres RO, McNally KL, Serraj R. Variation in root system architecture and drought response in rice (Oryza sativa): Phenotyping of the OryzaSNP panel in rainfed lowland fields. F Crop Res. 2011;120: 205–214. doi:https://doi.org/10.1016/j.fcr.2010.10.003

110. Martin M. Cutadapt removes adapter sequences from high-throughput sequencing reads. EMBnet.journal. 2011;17: 10. doi:10.14806/ej.17.1.200

111. Kim D, Salzberg SL. TopHat-Fusion: an algorithm for discovery of novel fusion transcripts. Genome Biol. 2011;12: R72. doi:10.1186/gb-2011-12-8-r72

112. Trapnell C, Williams BA, Pertea G, Mortazavi A, Kwan G, van Baren MJ, et al. Transcript assembly and quantification by RNA-Seq reveals unannotated transcripts and isoform switching during cell differentiation. Nat Biotechnol. 2010/05/02. 2010;28: 511–515. doi:10.1038/nbt.1621

113. Kærn M, Elston TC, Blake WJ, Collins JJ. Stochasticity in gene expression: from theories to phenotypes. Nat Rev Genet. 2005;6: 451–464. doi:10.1038/nrg1615

114. Schwabe A, Dobrzyński M, Rybakova K, Verschure P, Bruggeman FJ. Origins of Stochastic Intracellular Processes and Consequences for Cell-to-Cell Variability and Cellular Survival Strategies. In: Jameson D, Verma M, Westerhoff HVBT-M in E, editors. Methods in Systems Biology. Academic Press; 2011. pp. 597–625. doi:10.1016/B978-0-12-385118-5.00028-1

115. De Vos D, Bruggeman FJ, Westerhoff H V, Bakker BM. How Molecular Competition Influences Fluxes in Gene Expression Networks. PLoS One. 2011;6: e28494. Available: https://doi.org/10.1371/journal.pone.0028494

116. Shu X, Singh M, Karampudi NBR, Bridges DF, Kitazumi A, Wu VCH, et al. Xenobiotic Effects of Chlorine Dioxide to Escherichia coli O157:H7 on Non-host Tomato Environment Revealed by Transcriptional Network Modeling: Implications to Adaptation and Selection. Frontiers in Microbiology. 2020. p. 1122. Available: https://www.frontiersin.org/article/10.3389/fmicb.2020.01122

117. MacNeil LT, Walhout AJM. Gene regulatory networks and the role of robustness and stochasticity in the control of gene expression. Genome Research. 2011. doi:10.1101/gr.097378.109

118. Alonso JM, Stepanova AN, Leisse TJ, Kim CJ, Chen H, Shinn P, et al. Genome-wide insertional mutagenesis of Arabidopsis thaliana. Science (80-). 2003;301: 653–657. doi:10.1126/science.1086391

119. Harb A, Pereira A. Screening Arabidopsis Genotypes for Drought Stress Resistance. Plant Reverse Genetics Methods in Molecular Biology (Methods and Protocols). Totowa, NJ: Humana Press; 2011. pp. 191–198.

